# Comprehensive characterization of neutrophil genome topology

**DOI:** 10.1101/100198

**Authors:** Yina Zhu, Ke Gong, Matthew Denholtz, Vivek Chandra, Mark P. Kamps, Frank Alber, Cornelis Murre

## Abstract

Neutrophils are responsible for the first line of defense against invading pathogens. Their nuclei are uniquely structured as multiple lobes that establish a highly constrained nuclear environment. Here we found that neutrophil differentiation was not associated with large-scale changes in the number and sizes of topologically associating domains. However, neutrophil genomes were enriched for long-range genomic interactions that spanned multiple topologically associating domains. Population-based simulation of spherical and toroid genomes revealed declining radii of gyration for neutrophil chromosomes. We found that neutrophil genomes were highly enriched for heterochromatic genomic interactions across vast genomic distances, a process named super-contraction. Super-contraction involved genomic regions located in the heterochromatic compartment in both progenitors and neutrophils or genomic regions that switched from the euchromatic to the heterochromatic compartment during neutrophil differentiation. Super-contraction was accompanied by the repositioning of centromeres, pericentromeres and Long-Interspersed Nuclear Elements (LINEs) to the neutrophil nuclear lamina. We found that Lamin-B Receptor expression was required to attach centromeric and pericentromeric repeats but not LINE-1 elements to the lamina. Differentiating neutrophils also repositioned ribosomal DNA and mini-nucleoli to the lamina: a process that was closely associated with sharply reduced ribosomal RNA expression. We propose that large-scale chromatin reorganization involving super-contraction and recruitment of heterochromatin and nucleoli to the nuclear lamina facilitate the folding of the neutrophil genome into a confined geometry imposed by a multi–lobed nuclear architecture.

---

It is now well established that in interphase nuclei the chromatin fiber is folded into loops that are anchored by architectural proteins such as CTCF and members of the cohesion complex. Clusters of loops are organized as topologically associating domains (TADs). TADs, in turn, collectively associate with either the euchromatic (A) or heterochromatic (B) compartment. Finally, individual chromosomes fold into territories that only sporadically intermingle (Lieberman-Aiden et al. 2009; Rao et al. 2014; Jin et al. 2013; Dixon et al. 2012, 2015). An additional layer of chromatin architecture involves interactions between the chromatin fiber and nuclear structures (Bickmore and Van Steensel 2013). Particularly well-characterized nuclear structures are the nuclear lamina, Polycomb Group (PcG) bodies, transcription factories and nucleoli. Prominent among interactions involving the chromatin fiber and nuclear structures are the heterochromatin and the nuclear lamina (Towbin et al. 2013; Peric-Hupkes et al. 2010; Meuleman et al. 2013; Melcer and Meshorer 2010), a process mediated by the Lamin B receptor (LBR) and Lamin A/C (Solovei et al. 2013). Likewise, PcG bodies associate with H3K27me3- marked transcriptionally silent chromatin to form an intra- and inter-chromosomal network (Denholtz et al. 2013; Vieux-Rochas et al. 2015). Clusters of genes that are coordinately transcribed tend to cluster as transcription factories (Apostolou and Thanos 2008; de Wit et al. 2013; Li et al. 2012). Finally, ribosomal DNA (rDNA) repeat elements are arranged as nucleolar organizer regions that cluster when associated with the RNA polymerase I transcription machinery to form nucleoli, a nuclear body specialized in ribosomal RNA (rRNA) transcription, processing and assembly of ribosomes (Kalmárová et al. 2007; Németh and Längst 2011).

Neutrophils are terminally differentiated, short-lived circulating cells. Like other leukocytes, they are continuously generated from hematopoietic stem cells and develop through intermediate developmental stages. Neutrophils belong to a special class of immune cells called polymorphonuclear cells which also include basophils and eosinophils. Polymorphonuclear cells are unique in that their nuclei are ring-shaped or multi-lobulated, a feature that facilitates rapid migration through the endothelial lining of blood vessels and the interstitial spaces of tissues (Kolaczkowska and Kubes 2013). How the genomes of polymorphonuclear cells are organized into multiple lobes remains largely unknown. A critical component involves the LBR. Homozygous nonsense mutations in the *Lbr* gene result in an anomaly that is characterized by a failure to generate multilobular nuclei and an accumulation of immature neutrophils (Gaines et al. 2008; Verhagen et al. 2012). Hypolobulated neutrophils are deficient in their ability to migrate through interstitial spaces (Gaines et al. 2008).

Here we mapped architectural changes in genome organization during neutrophil differentiation. We found that the genome of neutrophils was characterized by an extensive loss of local genomic interactions (< 3 Mb) and a gain of long-range chromatin interactions (> 3 Mb) that spanned vast genomic distances. Population-based simulation of spherical and toroid genomes revealed declining radii of gyration for neutrophil chromosomes. We found that the neutrophil genome underwent widespread chromosome contraction, across vast genomic distances, a process termed super-contraction. Super-contraction involved genomic regions located in the B compartment in both progenitors and neutrophils or genomic regions that switched from the A to the B compartment during neutrophil differentiation. The differentiation of neutrophils was also accompanied by the repositioning of centromeres, pericentromeres, LINE-1 elements, rDNA and nucleoli from the nuclear interior to the nuclear lamina. We found that LBR expression was required to attach centromeric and pericentromeric repeats but not LINE-1 elements to the neutrophil nuclear lamina. We propose that super-contraction and recruitment of heterochromatin to the lamina facilitates the folding of neutrophil genomes into a confined geometry imposed by a multi-lobular nuclear structure. Furthermore, we suggest that rDNA localization regulates protein synthesis and neutrophil life span.

## Results

### A model system to study multi-lobed nuclear architecture

To examine how the neutrophil genome folds into a confined environment imposed by a multi-lobular nuclear structure we used a previously described cell line that permits the isolation of large numbers of homogeneous myeloid progenitors and multilobular cells. This system employs a pro-myeloid cell line named ECOMG, which was established by conditional immortalization of myeloid progenitors using an E2A/PBX1-estrogen receptor fusion protein (Sykes et al. 2003). As previously reported, ECOMG cells proliferated indefinitely in the presence of GM-CSF and estrogen but upon estrogen withdrawal readily differentiated into a homogeneous population of multilobular cells (> 98%) (Supplemental Fig. S1A). To compare the transcriptomes of *in vitro* differentiated neutrophils to murine bone marrow derived neutrophils, RNA was isolated from ECOMG progenitors (progenitors) and *in-vitro* differentiated neutrophils (neutrophils). The transcriptomes of both populations were analyzed using RNA-Seq and compared to the transcription signatures of murine bone marrow derived neutrophils (BMDN) (Wong et al. 2013). We identified 3192 transcripts that were expressed at higher abundance versus 3378 transcripts that were expressed at lower abundance in differentiated neutrophils versus progenitors (Supplemental Fig. S1B). Notably, we found that the transcription signatures of *in vitro* differentiated neutrophils closely correlated with primary neutrophils isolated from the bone marrow (Supplemental Fig. S1C). Gene Ontology analysis revealed that the transcription signature of *in vitro* differentiated neutrophils was in agreement with previous microarray assays: neutrophils activated genes in neutrophils were enriched adhesion, cytokine production, migration, phagocytosis and bacterial killing (Supplemental Fig. S1D,E) whereas neutrophil repressed genes primarily encoded for proteins associated with metabolism, RNA processing and translation (Supplemental Fig. S1D). Prominent amongst the changes in gene expression was the coordinate decline in transcript abundance of ribosomal proteins and RNA Polymerase I subunits (Supplemental Fig. S1E; right panel). Taken together, these data indicate that the transcription signature of *in vitro* differentiated neutrophils resembles that of primary neutrophils and that the differentiation of neutrophils is closely associated with a decline in the expression of genes associated with protein synthesis.

### Neutrophil genomes are enriched for long-range genomic interactions

As a first approach to determine how the topology of the neutrophil genome differs from that of mononuclear cells we used genome-wide chromosome conformation capture analysis (Hi-C) (Rao et al. 2014). We found that neutrophils and BMDN when compared to progenitors were depleted of genomic interactions that spanned less than 3 Mb but were enriched for genomic interactions separated by genomic distances larger than 3 Mb (Fig. 1A). We next constructed chromosome-wide contact maps for autosomes as well as X-chromosomes (Fig. 1B). We found that short-range genomic interactions (diagonal) were depleted whereas interactions across large genomic distance (off-diagonal) were enriched in neutrophils (Fig. 1B). The differences in short-range versus long-range interactions were even more pronounced for X-chromosomes (Fig. 1B). The differences were particularly apparent in the differential heat maps constructed by subtracting contact frequencies in progenitors from neutrophils (Fig. 1B; far right panels). The contact matrices revealed a similar distribution for TADs when comparing progenitors versus neutrophils (Supplemental Fig. S2A). The size and numbers of TADs were highly consistent for both cell types (Supplemental Fig. S2B). However, the average intrachromosomal contact probability as a function of genomic distance differed between progenitors and neutrophils. Specifically, we found that intra-TAD interactions and inter-TAD interactions involving adjacent TADs (< 3 Mb) were depleted whereas genomic interactions that span multiple TADs (> 3 Mb) were enriched for neutrophils (Fig. 1C). We next used principal components analysis to identify the A and B compartments. Genomic regions that displayed a continuum of either positive or negative PC1 values were defined as PC1 domains (PDs) (Lieberman-Aiden et al. 2009). Neutrophils versus progenitors displayed increased numbers of PDs that were overall smaller in size (Supplemental Fig. S2C). Specifically, we identified 488 PDs that switched from the A to the B compartment, whereas 342 PDs switched from the B to the A compartment (Supplemental Fig. S2D; Supplemental Table S1). The size of PDs that switched during the transition from progenitors to neutrophils was significantly smaller than that of average PDs (Supplemental Fig. S2E). We found that genomic interactions involving large genomic distances (> 3 Mb) for both compartment A and B were enriched in neutrophils (Fig. 1D). To quantify long-range versus local genomic interactions we calculated the distal ratio (percentage of contacts that were separated by > 3 Mb) for progenitors and neutrophils. We found that the distal ratio was significantly higher in neutrophils when compared to progenitors and the differences of the distal ratio were inversely correlated with differences in PC1 values (Fig. 1E). Specifically, genomic regions that were associated with declining PC1 values or switched PC1 values often displayed changes in interaction patterns: from short-range interactions in progenitors to long-range genomic interactions in neutrophils (Fig. 1E, Supplemental Table S2). In sum, these data indicate that neutrophil genomes are depleted for local genomic interactions (< 3 Mb) but enriched for long-range genomic interactions that span vast genomic distances (> 3 Mb).

**Figure 1.**
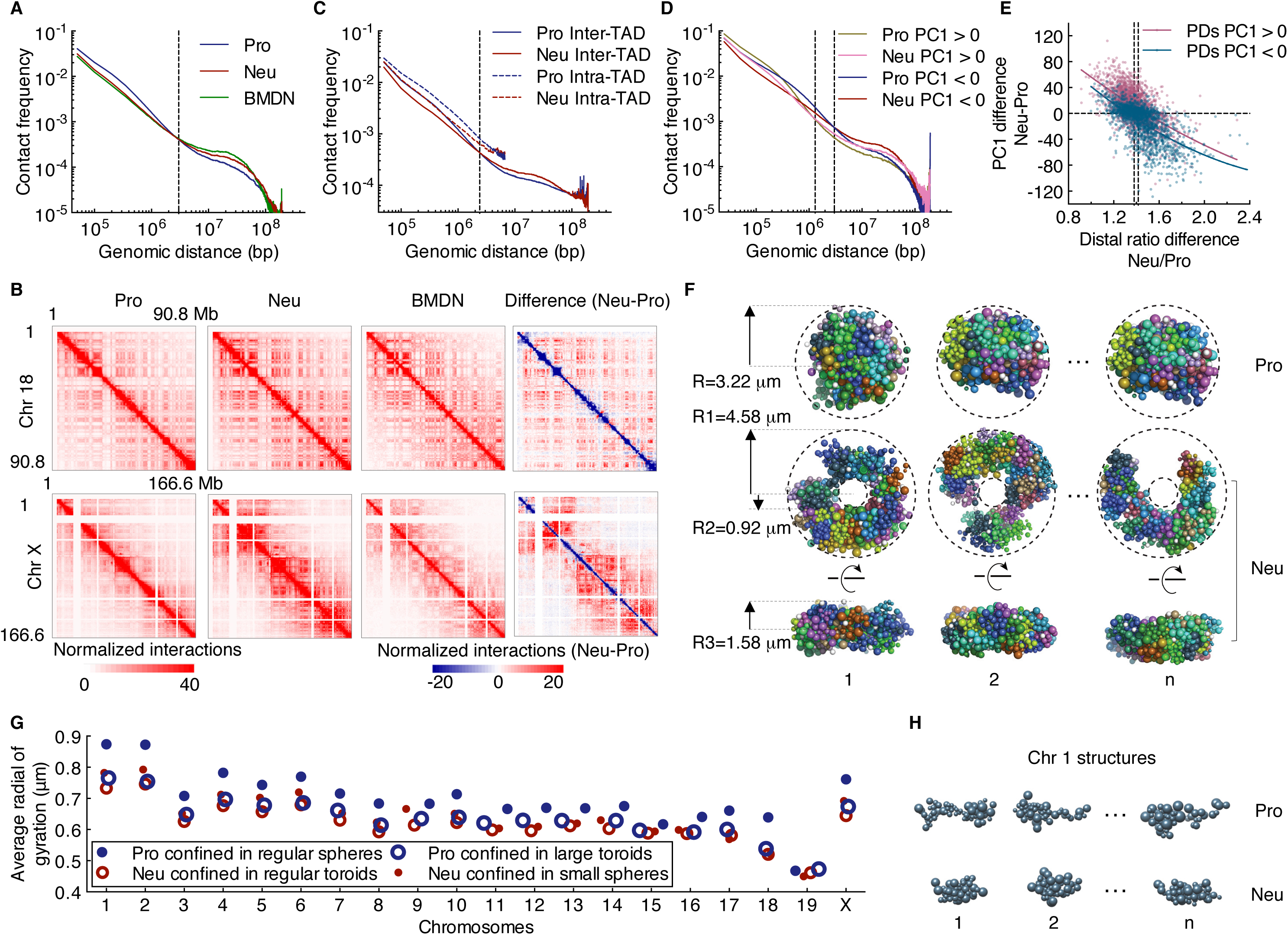
The differentiation of neutrophils is closely associated with large-scale changes in intradomain and inter-domain interactions. (A) Average contact probabilities as a function of genomic distance for genomic interactions within individual chromosomes at 50-kb resolution for progenitors (Pro), neutrophils (Neu) and bone marrow derived neutrophil (BMDN) genomes are shown. Dashed line indicates crossover for contact frequencies at 3Mb of gneomic separation. (B) Hi-C contact heatmaps for chromosome 18 (Chr 18) (top) and the X chromosome (Chr X) (bottom) at 500 kb resolution for progenitors, neutrophils and BMDN is shown. The intensity of each pixel represents normalized number of contacts between a pair of loci. Right panel indicates differential contact heatmaps for chromosome 18 and the X chromosomes. Subtraction of progenitor from neutrophil contact matrices shows changes in normalized number of contacts between the Hi-C samples. Red color represents enrichment whereas blue color represents depletion for normalized contact frequencies in neutrophils. (C) Intra-chromosomal contact frequency decay curves for intra- and inter-TAD interactions at 50-kb resolution for progenitors and neutrophils. (*D*) Intra-chromosome contact frequency decay curves for A and B compartments at 25-kb resolution for progenitor and neutrophil genomes are shown. Interactions were classified based on whether both end-points are located within the A or the B compartment. The crossover in contact frequencies between progenitor and neutrophil genomes is at 1.3 Mb and 3 Mb for A and B compartments, respectively (dashed lines). (*E*) Differences in distal ratio (fold change, Neu/Pro) versus the differences in PC1 values for neutrophil and progenitor genomes, for each PC1 continuous domain (PD) is indicated. Color intensity corresponds to the density of the data points. Solid lines indicated nonlinear fit curves. Genome-wide the distal ratio was significantly higher in neutrophils when compared to progenitors: a median distal ratio fold change of 1.38 for the A compartment versus 1.42 for B compartment (dashed lines). (*F*) Three representative structures for progenitor and neutrophil genomes were derived from the structure populations. Modeling was performed for genomes confined either as spherical or toroid nuclear volumes. Shown are top (middle panel) and side views (bottom panel) of modeled neutrophil nuclear structures. Dimensions and shape of the nuclear volumes are indicated in the panels and were based on experimental measurements (Supplemental Fig. S2F). (*G*) The average radius of gyration for each chromosome in neutrophil and progenitor genomes was computed for two different settings. The progenitor and neutrophil genomic structures were modeled using Hi-C reads derived from progenitor and neutrophil genomes folded into spherical or toroid nuclear structures. (*H*) Representative structures for chromosome 1 are shown for progenitors and neutrophils with a radius of gyration similar to the average value as indicated in resulting models.

### Simulating spherical and toroid genomes reveal decreased radii of gyration for neutrophil chromosomes

The unique nuclear shape confers a larger surface-to-volume ratio in neutrophils compared to progenitors (Fig. S2F). To determine how changes in nuclear shape affect genome compaction, we generated 3D models for mononuclear and multi-lobular genomes (Kalhor et al. 2012). Specifically, a population of genome structures was generated in which chromatin contacts were statistically consistent with the experimental Hi-C maps that were derived from progenitors and neutrophils (neutrophil: correlation 0.972, progenitor: correlation 0.970). Modeling was performed for genomes confined either in spherical or toroid nuclear shapes (Fig. 1F). We then computed the degree of compaction for each of the chromosomes by measuring the average radii of gyration (Kalhor et al. 2012). We observed substantial differences in the radii of gyration of chromosomes in the two cell types. In neutrophils the chromosomes were consistently more compacted with substantially lower radii of gyration when compared to progenitors (Fig. 1G,H). To investigate the role of the nuclear shape we also calculated, using Hi-C reads derived from progenitor cells, the radii of gyration of progenitor genomes confined in toroid nuclear shapes. Notably, progenitor chromosomes displayed smaller radii of gyration even though the nuclear volume remained unchanged (Fig. 1G). When changing neutrophil genomes from toroid to spherically nuclear shapes its chromosomes showed slightly increased radii of gyration, reflecting a more elongated chromosome shape (Fig. 1G). These observations suggest that differences in nuclear shape can impose distinct folding patterns of genomes: multi-lobular toroid-like nuclei tend to favor more compacted chromosome configurations.

### Super-contraction of neutrophil genomes

To determine how the increase in long-range genomic interactions as measured by Hi-C relates to spatial contraction we focused on a 13 Mb genomic region located on chromosome 17. This 13 Mb region includes the top five distal-favored PDs in neutrophil genomes as compared to progenitor genomes identified by distal ratio analysis (Supplemental Table S2). In neutrophils the 13 Mb region displayed significant enrichment for long-range genomic interactions (Fig. 2A,B). To visualize this region in single cells we performed 3D-FISH using fluorescently labeled BAC probes that spanned the 13 Mb region (Fig. 2B; red bars). We found that the majority of progenitors displayed three or four fluorescent foci versus one or two foci per allele for neutrophils, indicating that the increase in long-range genomic interactions as observed by Hi-C correlates with spatial contraction (Fig. 2C,D). Hereafter, we will refer to spatial contraction across large genomic distances as super-contraction. Taken together, these data indicate that the differentiation of neutrophils is closely associated with spatial contraction across large genomic distances, a process named super-contraction.

**Figure 2.**
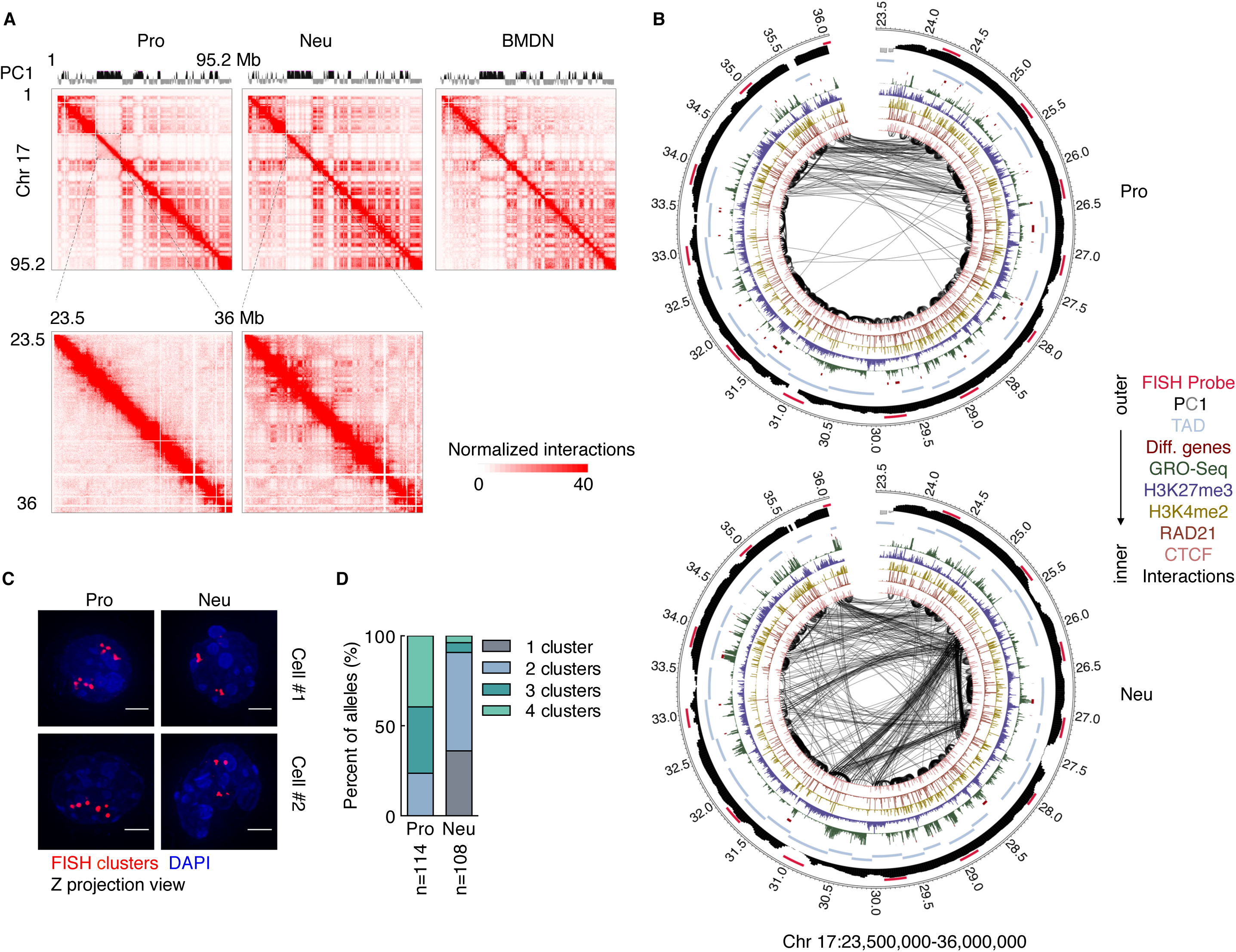
Genomes of differentiating neutrophils undergo large-scale spatial contraction. (*A*) Hi-C contact heatmap for chromosome 17 at 500-kb resolution (top) and a sub-genomic region for chromosome 17 at 50-kb resolution (bottom) for indicated cell types is shown. The intensity of each pixel represents the normalized number of contacts between a pair of loci. (*B*) Circos plots show genomic interactions across chromosome 17 derived from progenitor and neutrophil genomes. RAD21, CTCF, H3K4me2, H3K27me3, GRO-Seq, cell-type specific gene positions and PC1 values are shown. Probes used for 3D-FISH are indicated in red. Only significant interactions are shown: *P* < 0.001, binomial test. Thickness of the connecting lines reflects the significance of genomic interactions (-log P). Bin size, 50kb. Numbers around margins indicate genomic positions (in Mb). (*C*) 3D-FISH in progenitors and neutrophils using BAC probes spanning a 13-Mb region on chromosome 17 (*A*, *bottom*). Representative maximum intensity Z-projected images are shown. Original magnification, ×100. BAC probes, red; DAPI staining, blue. Scale bars, 2 μm. (*D*) Fractions of the number of clusters per allele were visualized using BAC probes for neutrophils and progenitors. n = number of alleles quantified for each sample.

### Epigenetic determinants that mark neutrophil differentiation

Previous observations indicated that developmental transitions are associated with the swiching of genes between A and B compartments (Lin et al. 2012; Dixon et al. 2015). To identify genes that switched compartments during neutrophil differentiation and how these changes relate to the activation of a neutrophil specific transcription signature we employed GRO-Seq. Overall nascent transcript levels correlated well with those observed for RNA-Seq (Supplemental Fig. S3A). Next we examined how during neutrophil differentiation changes in transcript levels relate to the switching of genes between nuclear compartments. We identified 65 genes that coordinately switched compartments and modulated levels of nascent transcription in differentiating neutrophils (Supplemental Fig. S3B). Out of a total of 1033 genes that in differentiating neutrophils were associated with increased nascent transcription 39 switched from compartment B to compartment A whereas 26 out of 2333 genes that displayed decreased nascent transcript levels switched from compartment A to compartment B (Supplemental Fig. S3B,F; Supplemental Table S3).

Given we found only a small proportion of genes that switched compartments exhibited coordinate changes in gene expression levels, we reasoned that other epigenetic mechanisms must exist that regulate a neutrophil specific transcription signature. Hence, we examined progenitors and neutrophils for the genome-wide deposition of 11 epigenetic marks (data not shown). Among these marks the deposition of H3K27me3 was particularly intriguing. Among the 1033 genes that were activated in neutrophils, we identified 222 genes associated with promoters that were enriched for H3K27me3 in progenitors but depleted of H3K27me3 in neutrophils (Supplemental Fig. S3D-F; Supplemental Table S3). Taken together, these data indicate that the activation of a neutrophil specific transcription signature is regulated at multiple levels that include changes in nuclear location but primarily involve depletion of H3K27me3 across promoters that are associated with genes that are activated in neutrophils.

### Differentiating neutrophils are enriched for long-range genomic interactions involving transcriptionally silent regions

To further evaluate the switching of genomic regions between compartments during neutrophil differentiation we segregated switched PDs as either transcriptionally active or transcriptionally silent domains as defined by the abundance of nascent transcription. We found that 82% of PDs that switched from compartment A to B were transcriptionally silent in both progenitors and neutrophils (Fig. 3A; Class I). Notably, Class I PDs were highly enriched for H3K27me3 and chromosomal contacts that spanned large genomic distances in neutrophils (Fig. 3B-D). Prominent among Class I PDs were the *Hox* loci. Multiple Class I PDs located in genomic regions that flank the *HoxD* locus, displayed a concerted decrease in intra-domain interactions and a concurrent increase in long-range genomic interactions in neutrophils (Fig. 3E,F). Previous studies have revealed that in ES cells genomic regions associated with the *HoxD* locus interact with other regions that are characterized by high abundance of H3K27me3 to generate a network of intra-and interchromosomal interactions across the A compartment (Denholtz et al. 2013; Vieux-Rochas et al. 2015). In contrast, we found that in neutrophils a network of genomic interactions, marked by H3K27me3, was associated with the B compartment. We next inspected progenitors and neutrophils for CTCF and RAD21 binding. CTCF occupancy remained unchanged during the transition from progenitors to neutrophils (Supplemental Fig. S4A). However we found that in differentiated neutrophils RAD21 occupancy was depleted across 4142 bound sites (Supplemental Fig. S4B). Inspection of significant interactions with endpoints in 5 kb regions surrounding CTCF peaks revealed similar interaction frequencies for both cell types (Supplemental Fig. S4C). When focused on interactions with endpoints across a 5 kb region surrounding differential RAD21 bound-sites we found that neutrophils displayed significantly lower interaction frequencies (Supplemental Fig. S4C). For example, within a 13 Mb region on chromosome 11, two domains that were depleted for RAD21 occupancy and maintained CTCF binding were depleted of intra-domain interactions but were enriched for intra-chromosomal genomic interactions that spanned vast genomic distances (Supplemental Fig. S5; upper panel, dotted purple boxes). Notably, both domains associated with this region were localized in the B compartment (Supplemental Fig. S5; upper panel, dotted purple boxes). Among the four classes of switched domains, class I PDs were most severely depleted of RAD21 occupancy (Supplemental Fig. S4D). Taken together, these data indicate that the differentiation of neutrophils is closely associated with *de novo* long-range chromosomal interactions involving genomic regions located in the B compartment in both progenitors and neutrophils or genomic regions that switched from the A to the B compartment that were enriched for H3K27me3 and depleted of RAD21 occupancy.

**Figure 3.**
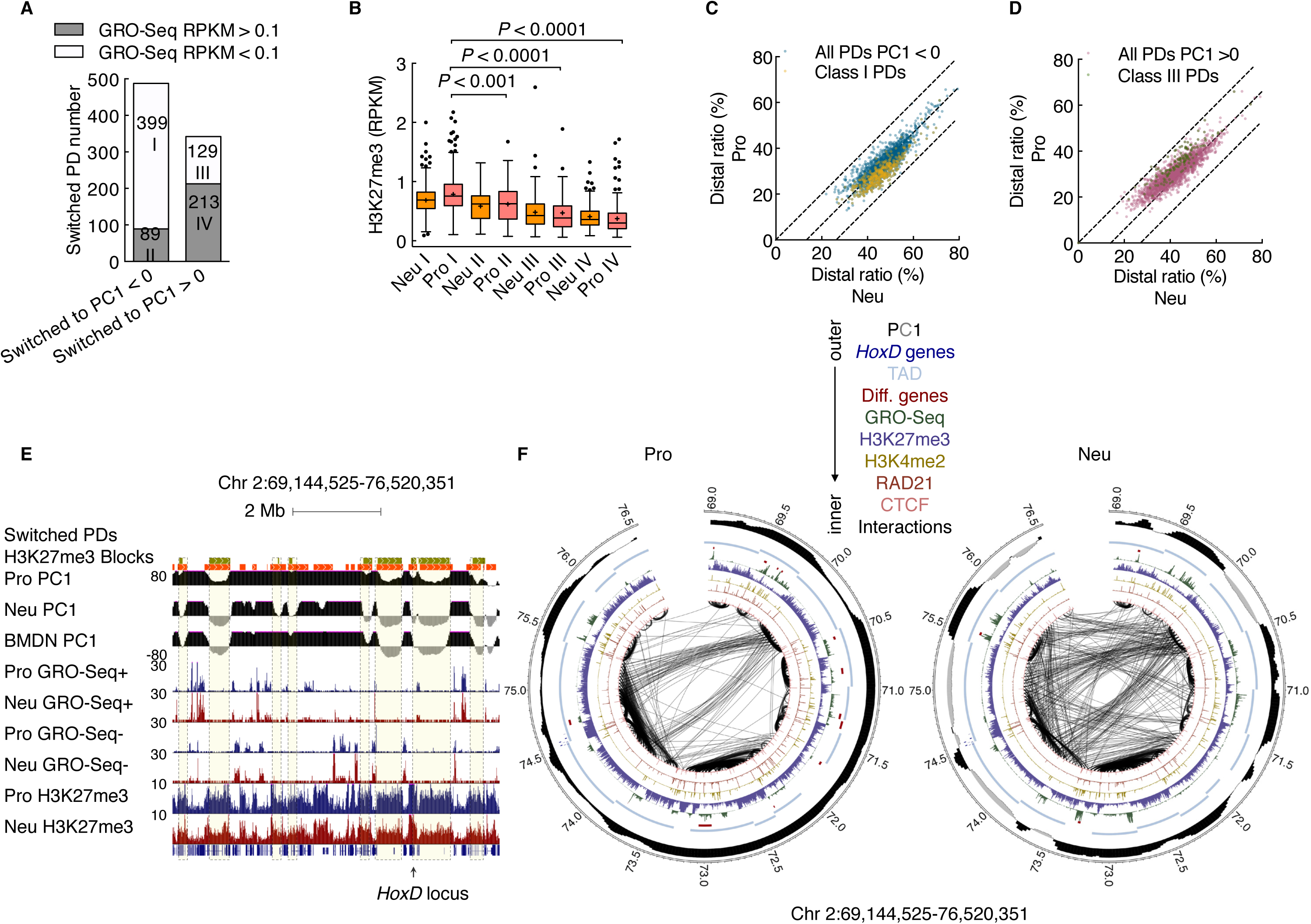
Differentiating neutrophils establish a network of genomic interactions involving transcriptionally silent regions. (*A*) Bar plot showing the numbers of four classes of PDs that switched compartments during neutrophil differentiation. The switched PDs were classified into two subcategories according to GRO-Seq transcript abundance: transcriptionally active, RPKM > 0.1 in either progenitors or neutrophils, or transcriptionally silent, RPKM < 0.1 in both progenitors and neutrophils. Numbers in bars indicate totals of PDs. Note that 82% of PDs that switched from the A to the B compartment in neutrophils (Class I) were transcriptionally silent in progenitors. (*B*) Box-and-whisker plot showing the abundance of H3K27me3 in four classes of PDs that switched compartments in progenitors and neutrophils. *P* value, Kruskal-Wallis test. (*C*) Scatter-plot comparing distal ratio of class I PDs and inactive PDs for neutrophil genomes. Color intensity corresponds to the density of the data points. (*D*) Scatter-plot comparing distal ratio of class III PDs and active PDs for neutrophil genomes. Color intensity corresponds to the density of the data points. (*E*) Genome browser snapshot of the *Hoxd* locus and flanking genomic regions. Deposition of H3K27me3-marked regions is shown for progenitors and neutrophils. Green filled boxes represent class I PDs. Orange filled boxes represent H3K27me3-marked regions. PC1 values as well as read densities for nascent RNA (GRO-Seq) and H3K27me3 are shown. (*F*) Circos plot showing genomic interactions within the same genomic region as in (*E*) for progenitor and neutrophil genomes are indicated. RAD21, CTCF, deposition of H3K4me2, H3K27me3, nascent transcripts abundance (GRO-Seq), cell-type specific gene positions and PC1 value are shown. Only significant interactions are shown: *P* < 0.001, binomial test. Thickness of the connecting lines reflects the significance of the interaction (-log *P*). Bin size, 50-kb. Numbers at the margins indicate genomic position (in Mb). Colors indicate CTCF, RAD21, H3K4me2, H3K27me3, GRO-Seq, cell-type specific genes, TADs, *HoxD* family, PC1 tracks and FISH probes (color key, up).

### Neutrophil genomes are enriched for inter-chromosomal interactions

To determine whether interchromosomal interactions were modulated during neutrophil differentiation we analyzed progenitors and neutrophils for inter-chromosomal contact frequencies (Kalhor et al. 2012). We found that inter-chromosomal contacts were highly enriched in neutrophil genomes when compared to progenitors (Fig. 4A,B). For individual chromosomes we found that centromere-proximal-ends (genomic regions located within ~20 Mb from chromosome ends) were most enriched for interchromosomal interactions in differentiated neutrophils (Fig. 4C,D). We next constructed genome-wide contact maps. We found that although the chromosomal interaction pattern appeared largely unchanged, all pairs of centromere-proximal-ends displayed enrichment for genomic contacts in neutrophils when compared to progenitors (Fig. 4E). A similar trend was observed for primary bone marrow derived neutrophils (Fig. 4E). To validate these findings in single cells we performed 3D-FISH using fluorescently labeled centromeric probes (Fig. 4F). Consistent with the Hi-C contact map we found that neutrophils displayed decreased numbers of centromeric fluorescent foci compared to progenitors, a pattern indicative of spatial clustering (Fig. 4G). Taken together these observations indicate that neutrophil differentiation is associated with enrichment for inter-chromosomal interactions that involve centromere-proximal-ends.

**Figure 4.**
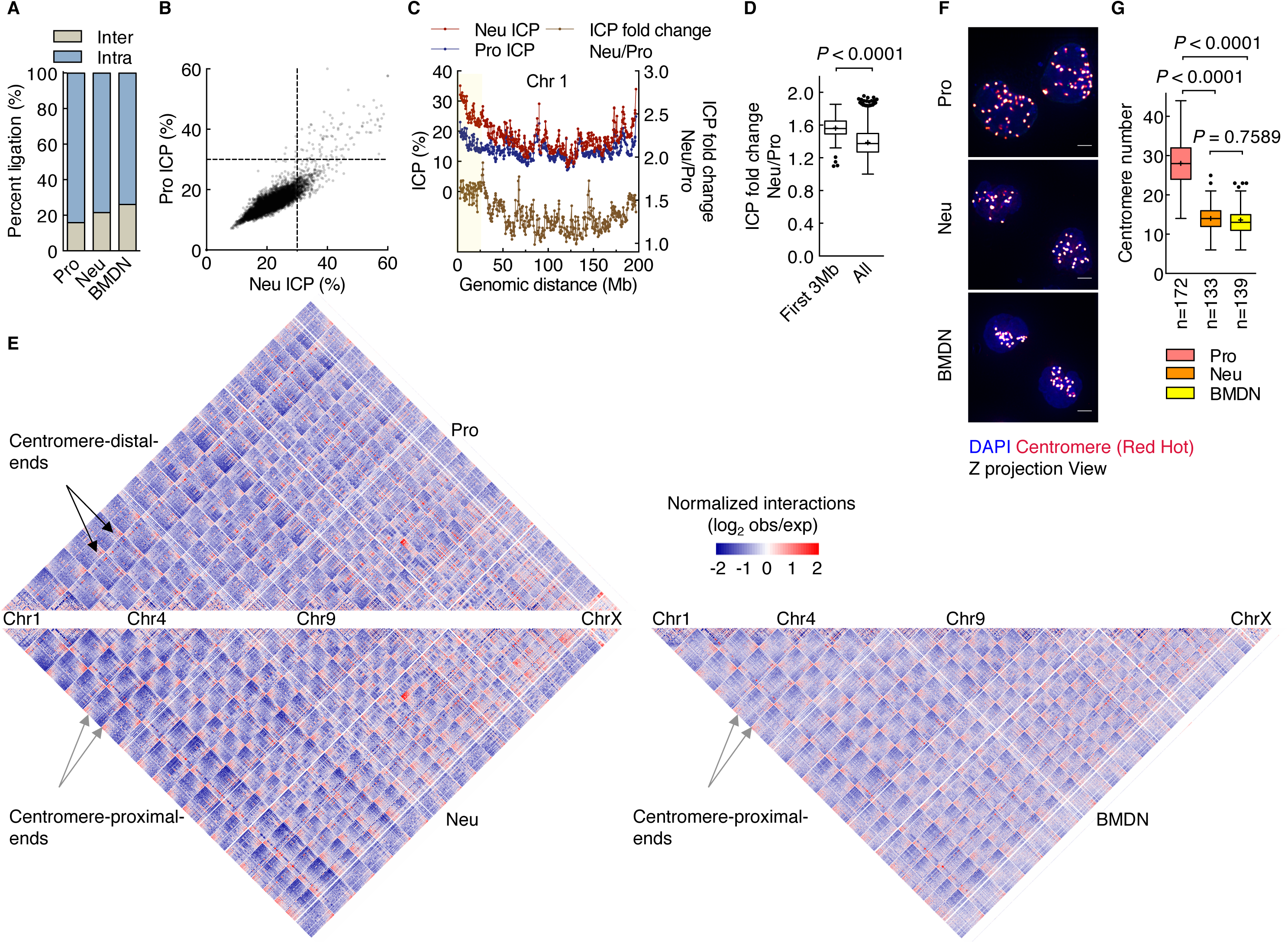
Differentiating neutrophils are enriched for interchromosomal interactions. (*A*) Neutrophil and BMDN genomes exhibit higher percentages of interchromosomal interactions than progenitor genomes. The fractions of intra- and interchromosomal contacts from progenitors, neutrophils and BMDN are indicated. (*B*) Scatter-plot comparing ICP values derived from neutrophil and progenitor genomes. ICP values were calculated as the ratio of interchromosomal interaction frequency versus the frequencies of all genomic interactions for each 500-kb interval. Color intensity corresponds to the density of data points. (*C*) Plots showing ICP values derived from progenitors and neutrophils (red and blue lines, left y-axis) and ICP differences (Neu/Pro) between the two cell types (yellow line, right y-axis) for chromosome 1 at 500kb resolution. (*D*) Box-and-whisker plot showing ICP fold change (Neu/Pro) comparison for centromeric proximal regions (< 3 Mb) to genome-wide average fold change values. *P* value, Mann-Whitney test. (*E*) Hi-C contact heatmap comparison between neutrophils (bottom left), BMDN (bottom right) and progenitors (top) at 500-kb resolution. Colors indicate the log ratio of observed interaction frequency to expected interaction frequency (obs/exp): blue, lower than expected; red, higher than expected. Black and gray arrows indicate centromere-distal-end-zone and centromere-proximal- end-zone, respectively. (*F*) 3D-FISH analysis of progenitors and neutrophils using a fluorescently labeled centromeric probe. Representative maximum intensity Z-projections of image stacks are shown. Original magnification, ×100. Red color indicates the intensity of centromere staining; DAPI staining, blue. Scale bars, 2 μm. (*G*) Box-and-whisker plot showing centromere numbers revealed using Volocity software. n = number of cells quantified for each sample. *P* value, one-way ANOVA test.

### LBR expression is required to attach centromeric and pericentromeric but not LINE-1 elements to the neutrophil lamina

To determine how centromeric and pericentromeric heterochromatin is positioned in progenitors and neutrophils, we used Immuno 3D-FISH. In progenitors, centromeres and major satellite repeats were localized both at the lamina and the nuclear interior whereas in neutrophils they primarily localized at the lamina (Fig. 5A,B,E). To determine how heterochromatic repeat elements were tethered at the neutrophil lamina we generated LBR-deficient progenitors using CRISPR-cas9 mediated deletion (Cong et al. 2013) (Supplemental Fig. S6A-C). *Lbr^l-^* progenitors were differentiated and examined for localization of pericentromeric and centromeric repeat elements (Fig. 5C-E). We found that centromeric regions in undifferentiated *Lbr^1-^* progenitors were numerous and primarily organized as small spherical compact bodies (Fig. 5C-E). In contrast, centromeres in *Lbr^l-^* neutrophils were fewer in number and localized as large clusters that were irregular in shape and located away from the lamina (Fig. 5C-E). Despite the striking repositioning of centromeric, pericentromeric DNA away from the lamina, we detected a rim of heterochromatin, reflected by DAPI staining, that remained associated with the lamina in differentiated *Lbr^1-^* neutrophils. To explore the possibility that LINEs were retained at the lamina of LBR-deficient neutrophils we performed 3D-FISH using a truncated LINE-1 element as probe. We found that in progenitors LINE-1 elements were dispersed throughout the nucleus but that in differentiated neutrophils they repositioned to the lamina (Fig. 6A-C). Notably and distinct from that observed for other heterochromatic repeat elements, a fraction of LINE-1 elements remained associated with the nuclear lamina whereas another fraction wrapped around chromocenters in LBR-deficient neutrophils (Fig. 6E). Thus, LBR expression is required to anchor centromeric, peri-centromeric but not LINE-1 elements to the neutrophil nuclear lamina.

**Figure 5.**
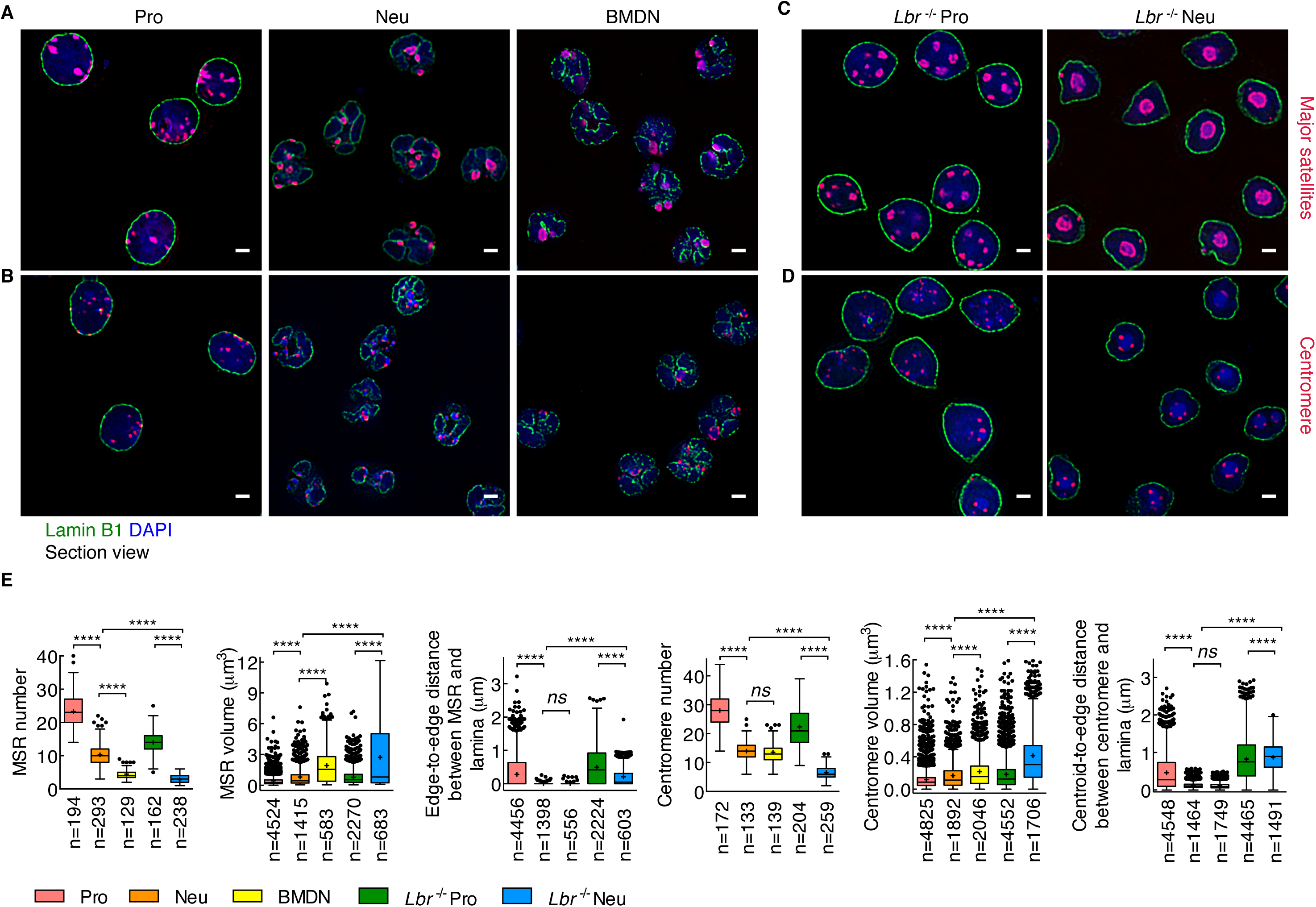
Pericentormeric and centromeric heterochromatin reposition to the lamina in differentiating neutrophils. (*A*) Pericentromeric heterochromatin in progenitors and neutrophils as well as BMDN were visualized using Immuno 3D-FISH. The lamina was stained using an antibody directed against Lamin B1. Pericentromeric DNA was identified using fluorescently labeled major satellite repeats (MSR) as a probe. Representative image section presented as digitally magnified images. Original magnification, ×100. FISH foci, red; Lamin B1, green; DAPI staining, blue. Scale bars, 2 μm. (*B*) Centromere in progenitors and neutrophils as well as BMDN were visualized as described in (*A*) but using a centromeric repeat element as a probe. (*C*) Major satellite repeat (MSR) elements in *Lbr*^-/-^ progenitors and *Lbr*^-/-^ neutrophils were visualized using immuno 3D-FISH as described in (A). (D) Centromeres in *Lbr*^-/-^ progenitors and *Lbr'^1-^* neutrophils were visualized using immuno 3D-FISH as described in (A). (E) Box-and-whisker plot showing the quantification of the number, volume and distance to the lamina of MSR and centromeres for progenitors, neutrophils, BMDN, *Lbr*^-/-^ progenitors and *Lbr*^-/-^ neutrophils using Volocity software. The distance of the MSR to the lamina was measured that separates the edge of the MSR from the edge of the lamina. The distance of the centromere to the lamina was measured using the centroid of centromere dots and the edge of the lamina. n = number of cells or FISH foci quantified for each sample. *P* value, one-way ANOVA test for numbers, Kruskal-Wallis test for volume or distance. ****, *P* < 0.0001; *ns*, not significant.

**Figure 6.**
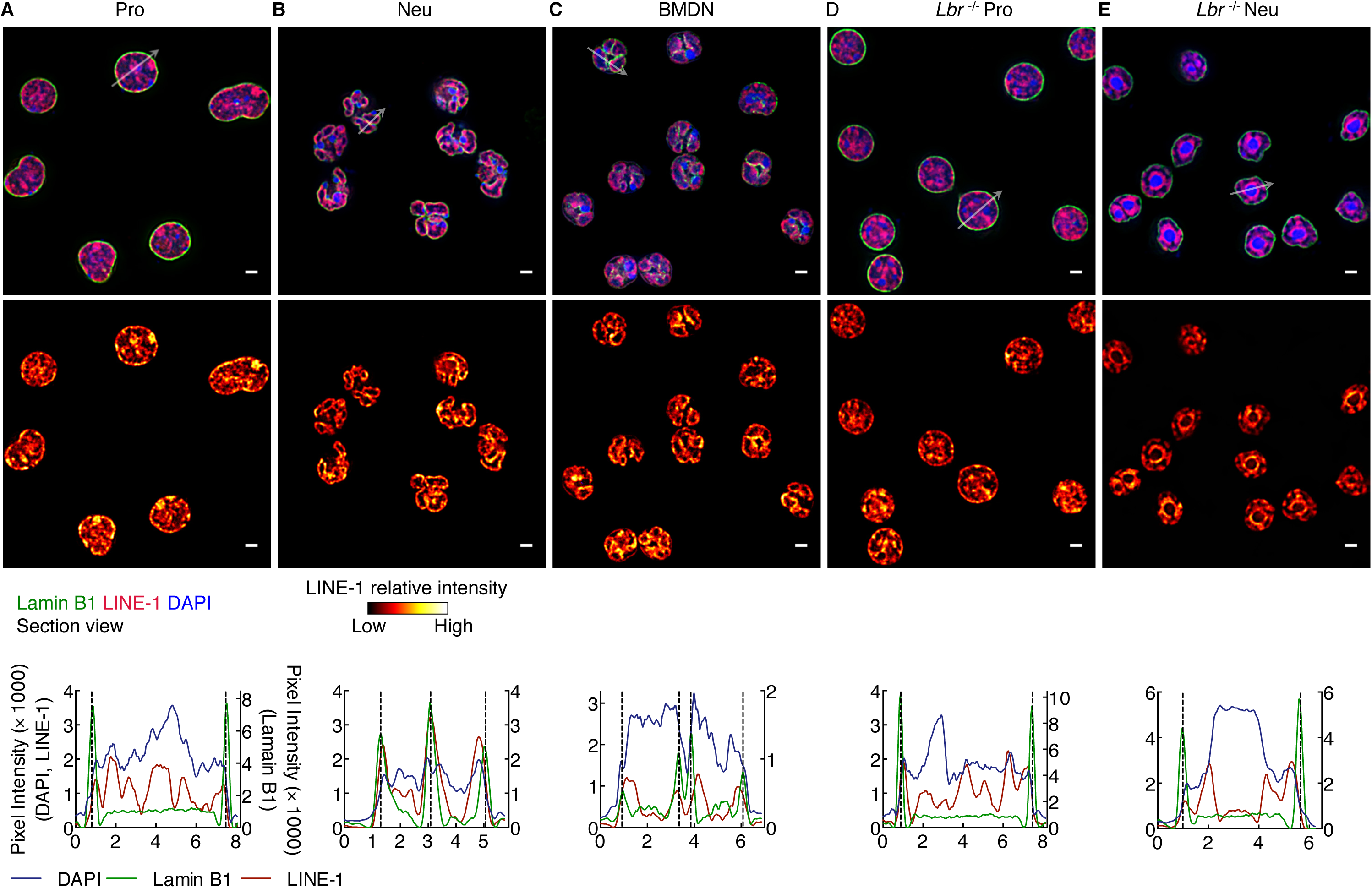
LINE-1 elements reposition during neutrophil differentiation. (*A-E*) The repositioning of LINE-1 elements to the nuclear lamina during neutrophil differentiation is independent of LBR expression. Top and middle panels: Immuno- FISH using antibodies directed against Lamin B1 antibody and a LINE-1 probe for progenitors (*A*), neutrophils (*B*), BMDN (*C*), *Lbr*^-/-^ progenitors (D) and *Lbr*^-/-^ neutrophils (*E*). Representative image sections are presented as digitally magnified images. Original magnification, ×100. Top: LINE-1, red; Lamin B1, green; DAPI staining, blue. Middle: red color scale indicates the intensity of FISH signal. Scale bars, 2 μm. White arrows indicate the diameter across the nucleus. Bottom: Radial distribution of fluorescence intensity signal for Lamin B1 (green line, right y-axis), LINE-1 and DAPI (red and blue lines, left y-axis).

### Repositioning of ribosomal DNA repeats in neutrophils

In addition to centromeres, pericentromeres and LINE-1 elements, rDNA constitutes yet another type of repeat element. To examine whether rDNA also repositions during neutrophil differentiation we used 3D-FISH. While clusters of rDNA in progenitors were readily detectable in the nuclear interior from progenitors, rDNA foci were far fewer in numbers and predominantly localized near the neutrophil nuclear lamina (Fig. 7A,B). To further characterize the nucleolus in differentiated neutrophils we examined progenitors and neutrophils for B23, a nucleolar protein that marks nucleoli. We found that during neutrophil differentiation nucleoli condensed in size and repositioned, in an LBR-dependent manner, to the nuclear lamina (Fig. 7C,D). To directly visualize the repositioning and disassembly of nucleoli in live cells we labeled nucleoli with B23-mCherry, heterochromatin with SUV39H1-GFP and the nuclear lamina with LBR-GFP. Briefly, progenitor cells were transduced with retroviruses expressing these fluorescent proteins and imaged in live progenitors and differentiated neutrophils. This analysis revealed that, consistent with the 3D-FISH analysis, the nucleolus condensed into small nuclear bodies that localized near the neutrophil lamina (Fig. 7E). To examine how the folding pattern of rDNA relates to the nucleolar structure we performed Immuno 3D-FISH (Fig. 7E). We found that in progenitors B23 was wrapped around arrays of rDNA but that this distinct structure disassembled in differentiated neutrophils (Fig. 7F). Since previous studies demonstrated that Lamin B1 expression maintains the functional plasticity of nucleoli we restored expression of Lamin B1 in neutrophils (Martin et al. 2009). We found that forced Lamin B1 expression was not sufficient to restore a nucleolar structure in differentiated neutrophils (Supplemental Fig. S7). Finally, to determine whether and how a change in nucleolar structure and nuclear location relates to rRNA abundance, we performed RNA-FISH. While ribosomal transcript abundance was high in progenitors, rRNA expression was barely detectable in differentiated neutrophils as well as in neutrophils isolated directly from bone marrow (Fig. 7G). Collectively, these data indicate that in differentiating neutrophils nucleoli reorganize and reposition to the nuclear lamina, a process that is closely associated with a halt in rRNA synthesis.

**Figure 7.**
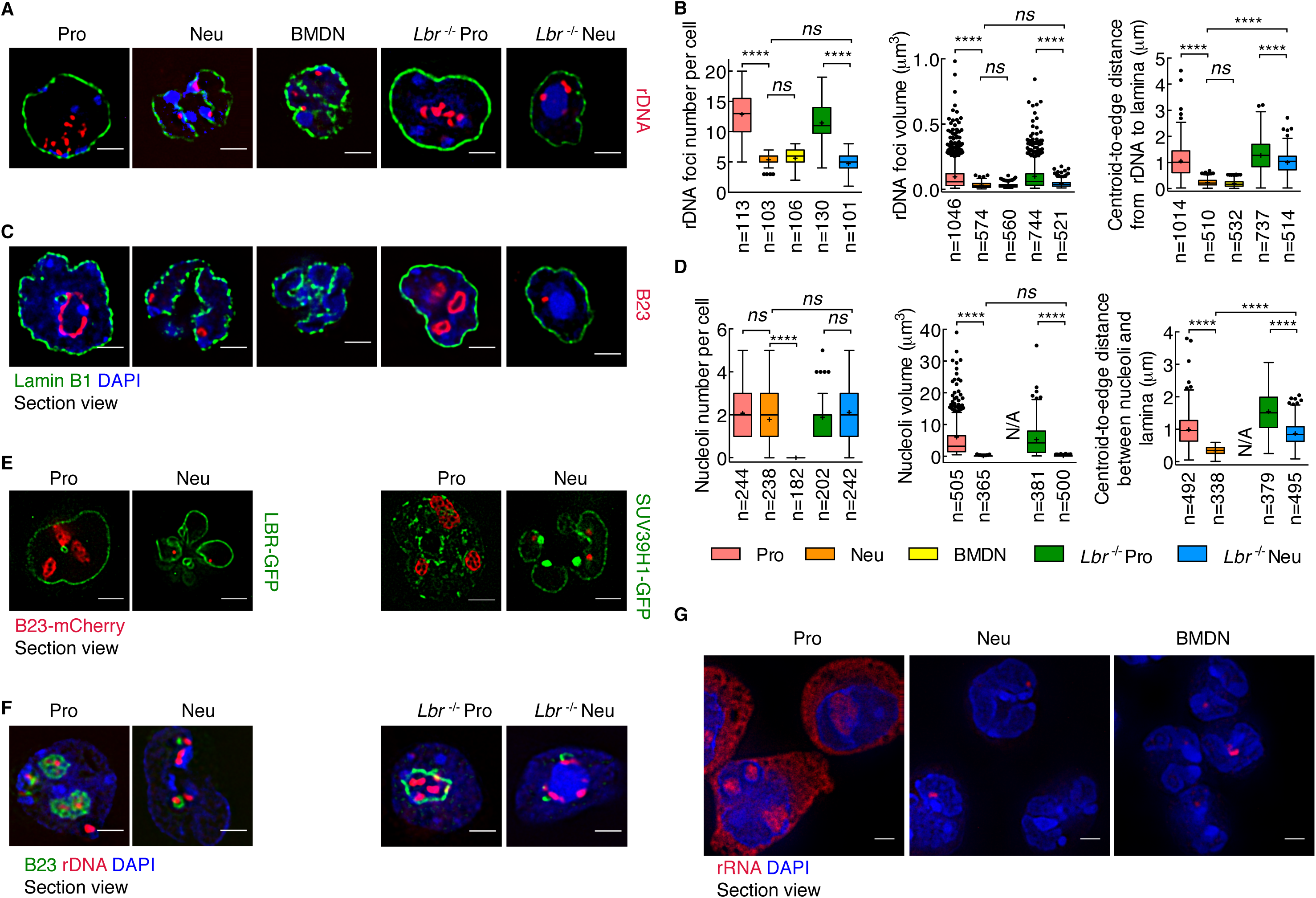
Spatial distribution of rDNA and nucleoli in differentiating neutrophils. (*A*) Ribosomal DNA repositions during neutrophil differentiation. Immuno-FISH using an antibody directed against Lamin B1 and a fluorescently labeled rDNA probe in progenitors, neutrophils, BMDN, *Lbr*^-/-^ progenitors and *Lbr*^-/-^ neutrophils. Representative image sections presented as digitally magnified images. Original magnification, χ100. rDNA, red; Lamin B1, green; DAPI staining, blue. Scale bars, 2 μm. (*B*) Box-and- whisker plot showing the quantification of the numbers, volumes and distances to the lamina of rDNA foci using Volocity software. The distance of rDNA foci to the lamina was measured between the centroid of dots and the edge of the lamina. n = number of cells or fluorescently labeled foci quantified for each sample. *P* value, one-way ANOVA test for numbers, Kruskal-Wallis test for volumes or distances. ****, *P* < 0.0001; *ns*, not significant. (C) Nucleoli reorganize during neutrophil differentiation. Immunofluorescence staining using an antibody directed against Lamin B1 and Nucleophosmin (B23) in progenitors, neutrophils, BMDN, *Lbr*^-/-^ progenitors and *Lbr*^-/-^ neutrophils. Representative image section presented as digitally magnified images. Original magnification, χ100. B23, red; Lamin B1, green; DAPI staining, blue. Scale bars, 2 μm. (D) Box-and-whisker plot showing the quantification of the numbers, volumes and distances to the lamina of nucleoli using Volocity software. The distance of nucleoli to the lamina was measured between the centroid of foci and the edge of the lamina. n = number of cells or nucleoli quantified for each sample. *P* value, one-way ANOVA test for number, Kruskal-Wallis test for volume or distance. ****, *P* < 0.0001; *ns*, not significant. (E) Live cell imaging in progenitors and neutrophils. To visualize changes in nucleolar structure in live cells, nucleoli were marked by B23- mCherry, the nuclear lamina was labeled using an LBR-GFP fusion protein and SUV39H1-GFP was generated to visualize the heterochromatin. ECOMG cells were transduced with vectors expressing LBR-GFP or SUV39H1-GFP and B23-mCherry. The transduced cells were differentiated into neutrophils. Representative image section shows the relative position of nucleoli to lamina and major satellites. Original magnification, *100. B23, red; LBR- or SUV39H1-GFP green; DAPI staining, blue. Scale bars, 2 μm. Note that in progenitors nucleoli were readily detectable using B23-mCherry as large distinct regions located in interior of the nucleus. Marked variation in size and shape, occasionally with very large and irregular shape was observed. Usually no more than 3 nucleoli per cell were detected. (F) Immuno-FISH using an antibody directed against B23 and fluorescently labeled rDNA probe in progenitors, neutrophils, *Lbr*^-/-^ progenitors and *Lbr*^-/-^ neutrophils. Representative image section shows that rDNA is bridging the mini-nucleolar structure to major satellites. Original magnification, ×100. rDNA, red; B23, green; DAPI staining, blue. Scale bars, 2 μm. *(G)* RNA-FISH using fluorescently labeled rDNA as a probe in progenitors, neutrophils and BMDN. Representative image sections are presented as digitally magnified images. Original magnification, ×100. rRNA, red; DAPI staining, blue. Scale bars, 2 μm. Note that the decrease in the nucleolar size in neutrophils is associated with a marked decline in nucleolar as well as cytoplasmic rRNA. The neutrophil mini-nucleolus body still maintains trace amounts of rRNA.

## Discussion

Among the wide spectrum of cells that comprise the immune system polymorphonuclear cells, including neutrophils, eosinophils and basophils, are unique in that their nuclei are organized into multiple lobes. Their special nuclear structure has raised the question as to how their genome architecture differs from that of mononuclear cells. We have utilized a previously developed *in vitro* model system of neutrophil differentiation and demonstrate that multi-lobular cells generated by this approach closely resemble neutrophils, in terms of transcription signatures and genome architecture, isolated from the murine bone marrow permitting a detailed study of how segmentation of nuclei into multiple lobes affects genome topology and how such changes related to neutrophil physiology. We found that the transition from a mononuclear to a multi-lobular nucleus was not associated with large-scale changes in the number and sizes of topologically associating domains. However, neutrophil differentiation was accompanied by a spectrum of alterations in genome topology, that included genomic interactions across vast genomic distances involving heterochromatic regions, switching of genomic regions from the euchromatic to the heterochromatic compartments and large-scale repositioning of repeat elements. Genomic regions that switch compartments and contract during developmental progression have been described previously. Prominent among these are loci encoding for regulators that control B cell fate, including EBF1 and FOXO1 (Lin et al. 2012). Likewise, antigen receptor loci switch nuclear location and undergo locus contraction in developing B cells (Jhunjhunwala et al. 2008; Lin et al. 2012). Super-contraction in neutrophils, however, is distinct since it involves transcriptionally silent regions and performs a structural rather a regulatory role. Super-contraction involves enrichment for genomic contacts involving genomic regions located in the B compartment or regions that switch from the A to the B compartment. The most striking epigenetic feature associated with super-contraction was the deposition of H3K27me3, marking pairs of interacting genomic regions that switched from the A to the B compartment. The repositioning from the A to the B compartment, however, was not associated with a domain-wise redistribution of H3K27me3. Hence it seems unlikely that H3K27me3 is the only player that promotes supercontraction. Rather we propose, as suggested by the simulation, that the remodeling of the genome into a toroid nuclear structure is imposed by physical forces that is facilitated by the deposition of H3K27me3 ultimately leading to large-scale chromosome contraction.

The changes in nuclear topology during neutrophil differentiation also involve the repositioning of genomic repeats including centromeres, pericentromeres and LINE-1 elements. LINE-1 elements are scattered across the genome, enriched in the B compartment and reposition at a global scale during neutrophil differentiation. A role for a subset of LINE-1 elements in structuring the genome has previously been proposed for X-chromosome inactivation and chromosome contraction (Chow et al. 2010; Corbel et al. 2013; Hall et al. 2014). Along the same line we suggest that during neutrophil differentiation LINE-1 elements promote chromosome contraction by sequestering large portions of the neutrophil genome to the lamina.

### Lamin B Receptor expression and the generation of multi-lobular nuclei

Finally, as previously observed and confirmed here, the generation of multilobular nuclei requires LBR expression neutrophils (Gaines et al. 2008; Verhagen et al. 2012). Additinally, we demonstrate here that LBR expression is essential to sequester pericentromeric heterochromain to the lamina but it is not essential for heterochromatic clustering in differentiating neutrophils. The mechanism by which LBR promotes a multi-lobular nuclear shape remains to be determined but can now be addressed experimentally using conditionally induced LBR-deficient ECOMG cells. Previous studies and the observations described here demonstrate that LBR expression in neutrophils is essential for the generation of multilobular nuclear structures and to sequester pericentromeric heterochromatin to the nuclear lamina. We note that neutrophils display substantial differences in nuclear morphology across species. The majority of neutrophils isolated from invertebrates, as well as several reptiles (snakes and turtles), named heterophils, are mononuclear in shape. It will be of interest to determine whether differences in LBR expression between heterophils and vertebrates multilobular cells underpin the distinctive features associated with neutrophil genomes across the animal kingdom.

### Repositioning of ribosomal DNA during neutrophil differentiation

During neutrophil differentiation, nucleoli mediate some of the most striking changes in nuclear architecture. We found that during neutrophil differentiation rDNA was sequestered at the lamina in a heterochromatic environment and associated with a virtual absence of rRNA expression. The decline in rRNA abundance may have implications for neutrophil physiology. It is well established that at least a subset of neutrophils are short-lived cells with a half-life in peripheral blood that is less than several hours. Although still to be proven it may very well be that the life span of neutrophils is controlled, at least in part, by rRNA abundance. We note that likewise other terminally differentiated cells, including erythrocytes, platelets, keratinocytes and lens fibre cells are associated with low ribosome numbers and it will be of interest to determine whether the repositioning of rDNA is also associated with decrease protein synthesis and life span in these cell types.

### Conclusion

In sum, we propose that the large-scale changes in nuclear topology established during neutrophil differentiation are orchestrated by both physical and molecular mechanisms that involve changes in geometric confinement imposed by a multi-lobular nuclear structure as well as enrichement for intrachromosomal interactions across vast genomic distances and the repositioning of repeat elements from the nuclear interior to the lamina.

## Materials and methods

### Cell culture and plasmids

Cells were maintained in a 37°C humidified incubator in the presence of 5% CO2. HEK-293T cells were cultured in DMEM (Invitrogen, 11995-073) supplemented with 10% fetal bovine serum and penicillin-streptomycin-glutamine (Invitrogen, 10378-016). GM-CSF-dependent ECOMG cells were grown in RPMI 1640 (Invitrogen, 11875-093) with 10% FBS and penicillin-streptomycin-glutamine, by the addition of 1:100 conditioned media (about 10 ng/mL GM-CSF) isolated from a B16 melanoma cell line stably transfected with a murine GM-CSF construct. β-estradiol (Sigma, E2758), where applicable, was added to the medium at a final concentration of 1 μM from a 10,000 × stock in 100% ethanol. Differentiation of ECOMG cells into granulocytes was induced by withdrawal β-estradiol as previously described (Sykes et al. 2003). CD11b^+^Ly6G^+^ was used as a marker to validate granulocytic differentiation at day 5 (data not shown).

In order to ectopically express Lamin B1-GFP, LBR-GFP, SUV39H1-GFP and B23-mCherry, their coding sequences were amplified from ECOMG cell cDNA. Via BamHI sites, the PCR products were ligated into the multiple cloning site of pMys-IRES-TAC vector. Retroviral supernatant was obtained through HEK-293T transfection using calcium phosphate precipitation and vectors encoding for Lamin B1-GFP, LBR-GFP, SUV39H1-GFP and B23-mCherry in conjunction with the packaging plasmid pCL- Eco. Two days after infection, samples underwent enrichment for infected cells through the use of antihuman CD25 magnetic beads (Miltenyi biotech) and an AutoMACS separator (Miltenyi biotech). Plasmids and maps will be provided upon request.

### Population-based structure modeling

We generated models of 3D genome structures using our recently introduced population-based modeling approach, which constructs a population of three-dimensional genome structures derived from and fully consistent with the Hi-C data (Kalhor et al. 2012; Tjong et al. 2016). By embedding the genome structural model in 3D space and applying additional spatial constraints (e.g. all chromosomes must lie within the nuclear volume, and no two chromosome domains can overlap), it is possible to deconvolute the ensemble-based Hi-C data into a set of plausible structural states. The data pre-processing, normalization and modeling were described previously (Kalhor et al. 2012; Tjong et al. 2016). Chromatin is represented at the level of macrodomains at about 3-Mb resolution, which were inferred from the Hi-C data.

### Accession number

The Hi-C, Gro-Seq, RNA-Seq, ChIP-Seq and MeDIP-Seq data were deposited at the NCBI Gene Expression Omnibus (GEO) database and are accessible through GEO (GSE93127).

## Acknowledgments

Sequencing was performed at the IGM Genomics Center, University of California, San Diego, La Jolla, CA, supported by grant # P30CA023100. Imaging was performed at the microscopy core of the School of Medicine, University of California, San Diego, La Jolla, CA, supported by grant CA23100 and NS047101. Y.Z. was supported by a Cancer Research Institute postdoctoral fellowship. F.A. was supported by the NSF (CAREER 1150287) and the Arnold and Mabel Beckman Foundation. C.M. was supported by the NIH (DK107977, AI00880, AI09599 and AI102853).

## Supplemental Material

**Supplemental Figure S1.**
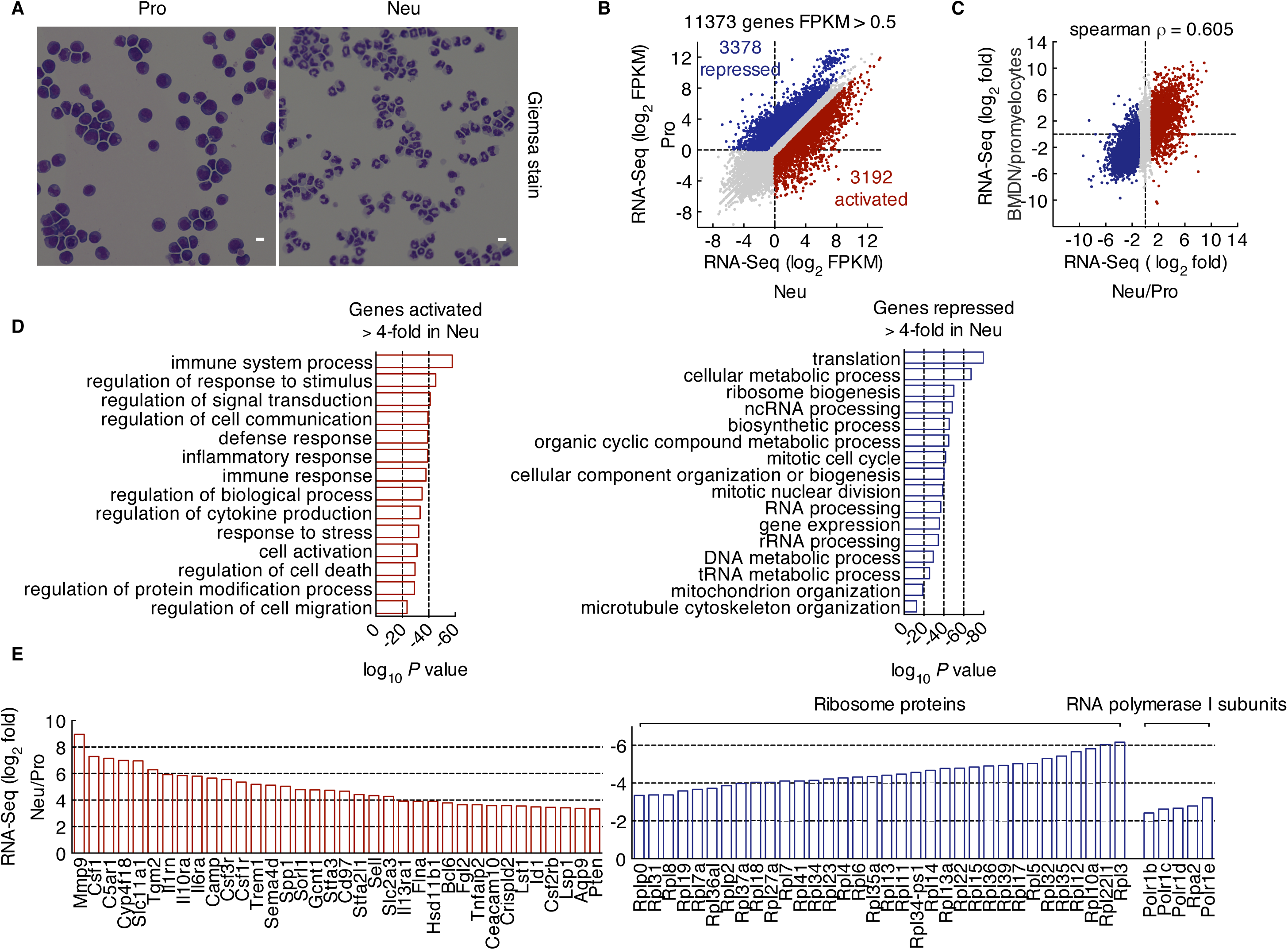
Transcription signatures of progenitors and *in vitro* differentiated neutrophils. (*A*) Morphological changes of ECOMG derived neutrophils after five days of cell culture in the absence of estrogen. Scale bars, 10 μm. (*B*) Scatter-plot comparing RNA-Seq profiles of ECOMG cells cultured with or without estrogen for 5 days. Colors indicate transcript abundance in neutrophils when compared to progenitors (less abundant-blue or more abundant-red) (difference of over 2-fold for each, FPKM > 0.5). (*C*) Scatter-plot comparing differences in RNA transcript abundance (log_2_ fold) for *in-vitro* differentiated neutrophils and progenitors plotted against the difference in RNA transcript abundance (log_2_ fold) for bone marrow derived neutrophils (BMDN) and promyelocytes. Spearman correlation ρ = 0.605. (*D*) Top gene ontology terms (via the DAVID bioinformatics database) enriched for genes activated (left) repressed (right) by a factor of more than 4-fold in neutrophils when compared to progenitors. (*E*) Selected group of genes that were repressed or activated during the differentiation of progenitor cells to neutrophils. Data are from one experiment.

**Supplemental Figure S2.**
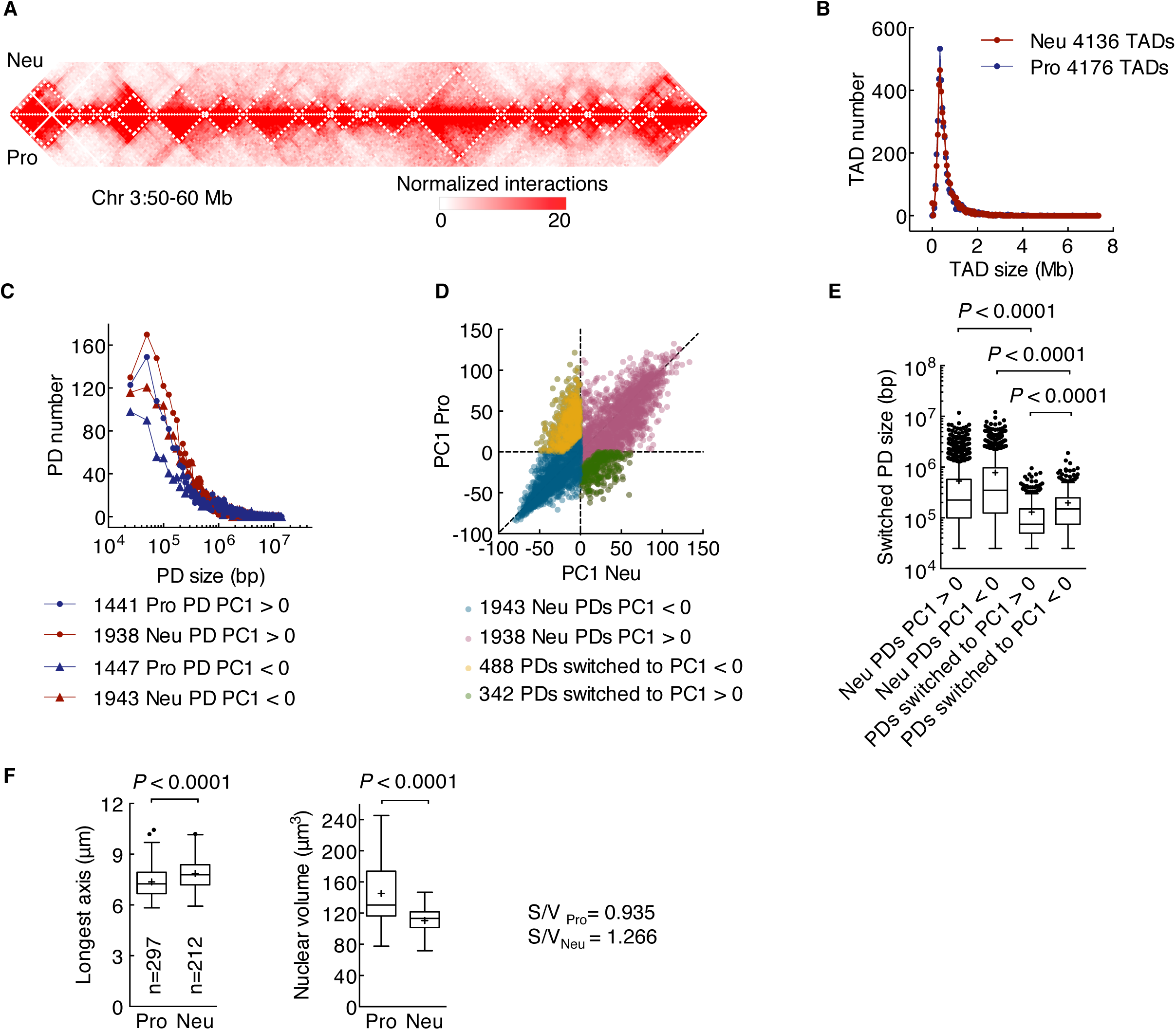
The organization of topologically associating domains and PC1 continuous domains in progenitor and neutrophil genomes. (*A*) Hi-C contact heatmap comparison between neutrophils (top) versus progenitors (bottom) at 50-kb resolution for a 10 Mb regions (Chr 3: 50,000,000-60,000,000). TADs were identified using TopDom domain calling method at 50-kb resolution using normalized matrices. The white dashed boxes indicate the shape of TADs. (*B*) Distribution of TAD size for progenitor and neutrophil genomes. TopDom resulted in 4176 TADs for progenitors, and 4136 TADs for neutrophils with a median size of 450 kb. (*C*) Distribution of PDs size across the genome, as defined by continuous regions of positive or negative PC1 values, for the A and the B compartments. (D) PC1 values associated with neutrophil and progenitor genomes. Signals away from the diagonal indicate PDs that switched nuclear compartments during neutrophil differentiation. (*E*) Box-and-whisker plot comparing the size of PDs. *P* value, Kruskal-Wallis test. (*F*) Box-and-whisker plot showing nuclear volume and longest axis of progenitors and neutrophils. Using Volocity 3D software, the volume and longest axis of DAPI stained nuclei of individual cells was determined by measuring the calibrated number of voxels identified within a given nucleus. Surface to volume ratios (S/V) are indicated. n = number of cells quantified for each sample. *P* value, Student's t test. Data are from one experiment.

**Supplemental Figure S3.**
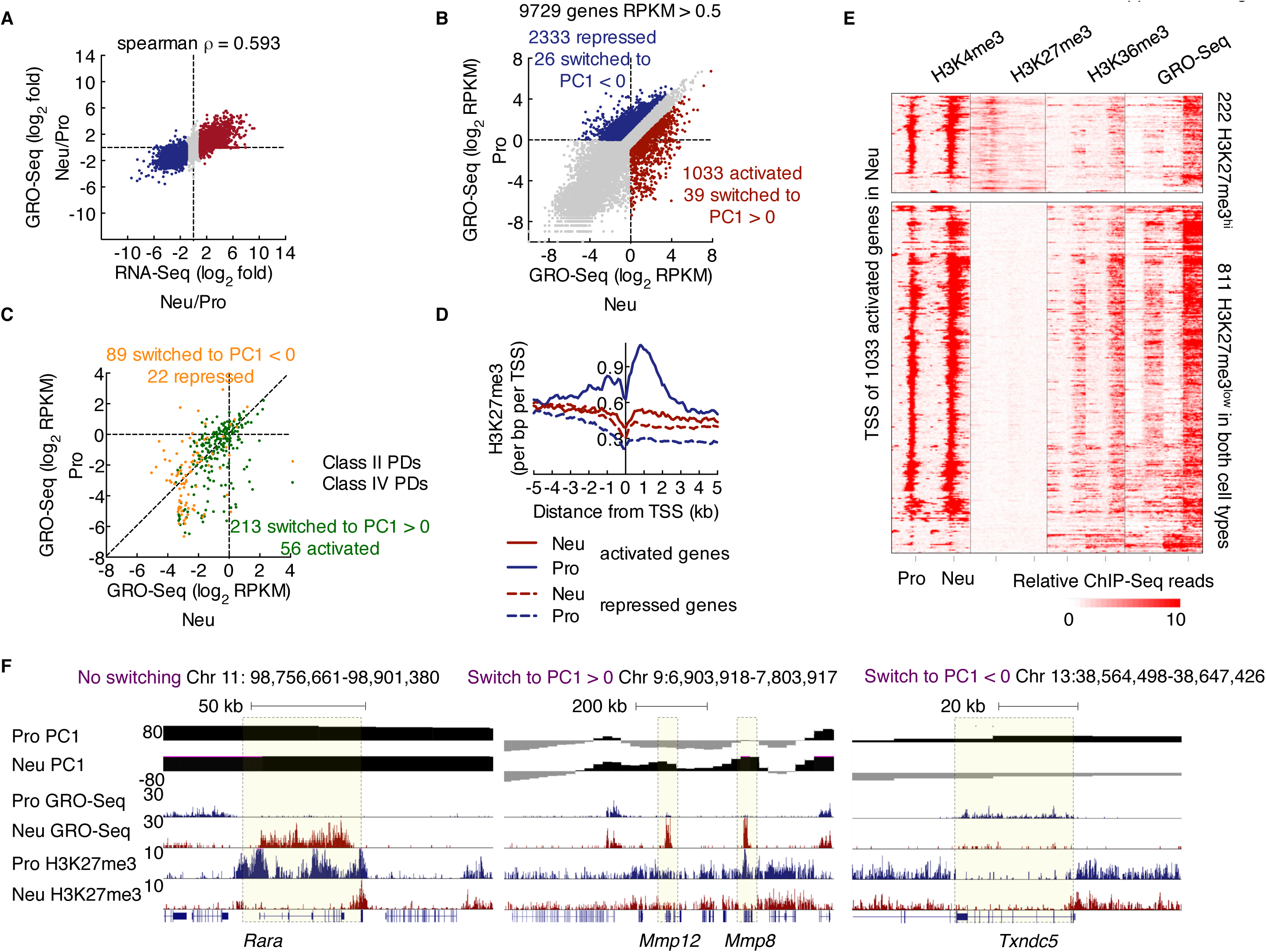
Transcription signatures and epigenetic marks associated with neutrophil differentiation. (*A*) Scatter-plot comparing the difference in nascent RNA transcript abundance (log_2_ fold; GRO-Seq) in neutrophils and progenitors versus the difference in steady-state mRNA abundance (log_2_ fold; RNA-Seq) in neutrophils and progenitors. Colors indicate transcripts levels in neutrophils than in progenitors as measured by RNA-Seq reads (less abundant-blue or more abundant-red) (difference of over 2-fold for each, FPKM > 0.5). Spearman correlation ρ = 0.593. (*B*) Scatter-plot comparing GRO-Seq associated with progenitors and neutrophils (difference of over 2-fold for each, RPKM > 0.5). Numbers in parenthesis indicate number of genes. Among the differentially expressed genes, 39 switched from the B to the A compartment; 26 switched from the A to the B compartment upon differentiating into neutrophils. (*C*) Scatter-plot comparing GRO-Seq profiles of switched PDs that were transcriptionally active (GRO-Seq RPKM > 0.1) from progenitors and neutrophils. Orange indicates class II PDs that switched from the A to the B compartment in neutrophils; green refers to class IV PDs switched from the B to the A compartment in differentiating neutrophils. Numbers in parenthesis indicate number of PDs. (*D*) Average H3K27me3 ChIP-Seq tag coverage in neutrophils and progenitors, plotted as a function of genomic distance from the TSS (± 5 kb), gated on 1033 neutrophil activated genes or 2333 neutrophil repressed genes. (*E*) Heatmap of ChIP-Seq data shows the distribution of H3K4me3, H3K27me3, H3K36me3 and GRO-Seq reads in progenitors and neutrophils gated on a window of ± 5 kb across the TSS of 1033 neutrophil activated genes. 222 TSS were associated with high levels of H3K27me3 in progenitors; 811 TSS were associated with low levels of H3K27me3 in both progenitors and neutrophils. Read density is displayed for a 10-kb window and color-scale intensities are shown in reads per million mapped reads per base pair. (*F*) Genome browser snapshots of (left) *Rara* locus (Chr 11: 98,756,661-98,901,380), (middle) *Mmp8* and *Mmp12* locus (Chr 9:6,903,918-7,803,917), (right) *Txndc5* locus (Chr 13:38,564,498-38,647,426), showing PC1 values for neutrophils and progenitors in a region surrounding this locus (top two rows), as well as read densities for nascent RNA (GRO-Seq) and H3K27me3 (bottom four rows). Light yellow shadows highlight the gene locus. Data are pooled from two independent experiments (GRO-Seq) or from one experiment representative of two independent experiments (H3K27me3 ChIP-Seq).

**Supplemental Figure S4.**
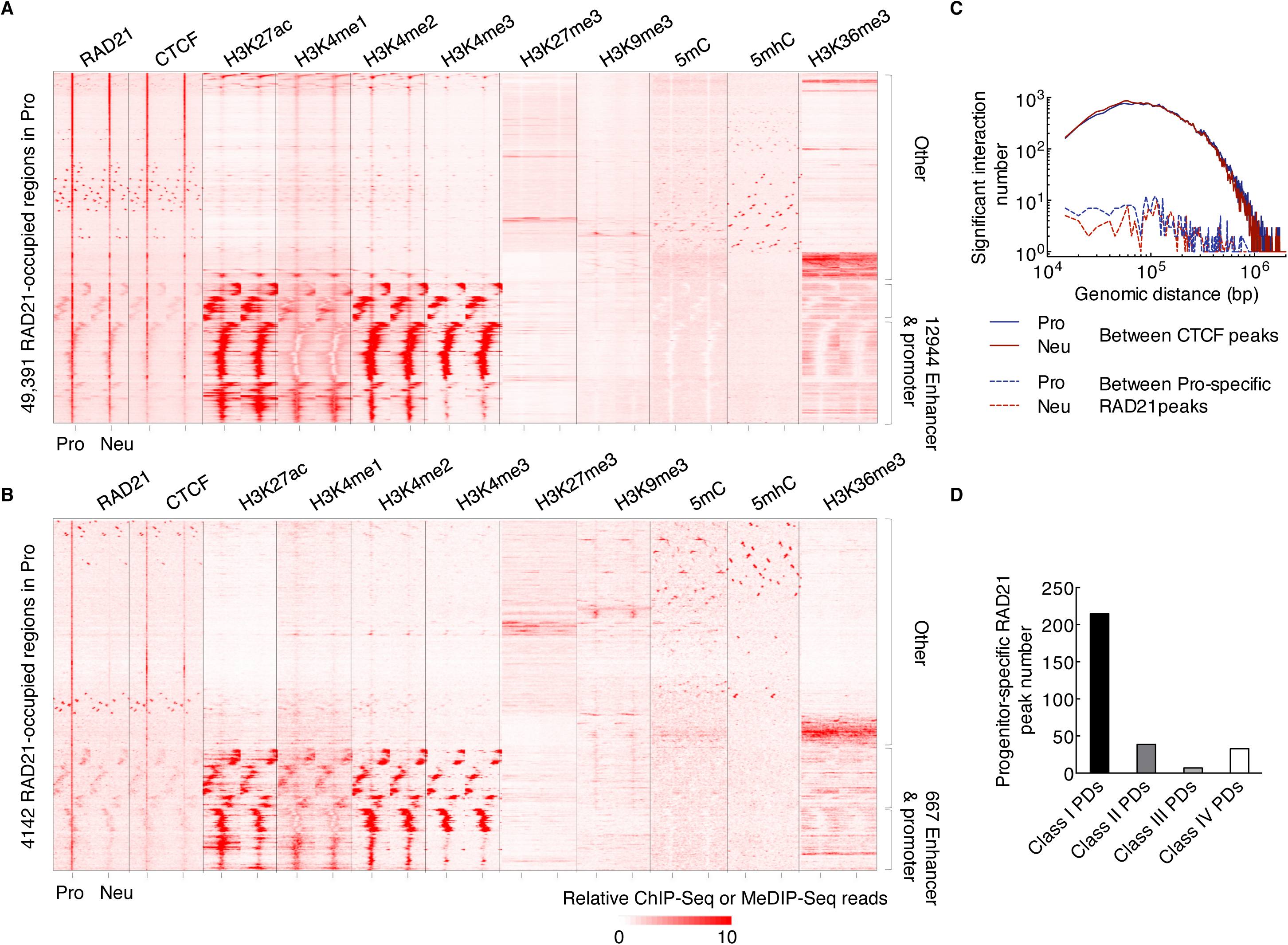
Genome-wide RAD21 occupancy in progenitors and neutrophils. (*A*) Heatmap of ChIP-Seq and MeDIP-Seq data showing the distribution of CTCF and RAD21 occupancy as well as the deposition of H3K27ac, H3K4me1, H3K4me2, H3K4me3, H3K27me3, H3K9me3, 5mC, 5mhC and H3K36me3 in progenitors and neutrophils, for a window of ± 5 kb across RAD21-bound sites that were gated on a total of 49,391 RAD21 bound sites identified in progenitors. The data for progenitors indicate that ~26% of cohesin-occupied sites involve enhancers and promoters. Read density is displayed for a 10-kb window and color-scale intensities are shown in reads per million mapped reads per base pair. (*B*) Heatmap of ChIP-Seq and MeDIP-Seq data showing the distribution of CTCF and RAD21 occupancy as well as the deposition of H3K27ac, H3K4me1, H3K4me2, H3K4me3, H3K27me3, H3K9me3, 5mC, 5mhC and H3K36me3 in progenitors and neutrophils, for a ± 5 kb window associated with 4142 progenitor-specific RAD21 peaks. The data for progenitors indicate that ~16% of RAD21-bound sites involved enhancers and promoters, ~17% of RAD21-bound sites were located close to enhancers and promoters and ~69% of bound sites were not associated with either enhancers or promoters. Read density is displayed for a 10-kb window and color-scale intensities are shown in reads per million mapped reads per base pair. (*C*) Genomic distance versus the number of significant interactions was plotted for CTCF versus CTCF interactions for pairs of CTCF-bound sites as well as pairs of progenitor specific RAD21-bound sites in progenitors and neutrophils. (*D*) Progenitor-specific RAD21 peak distribution among the four classes of switched PDs. Data are from one experiment.

**Supplemental Figure S5.**
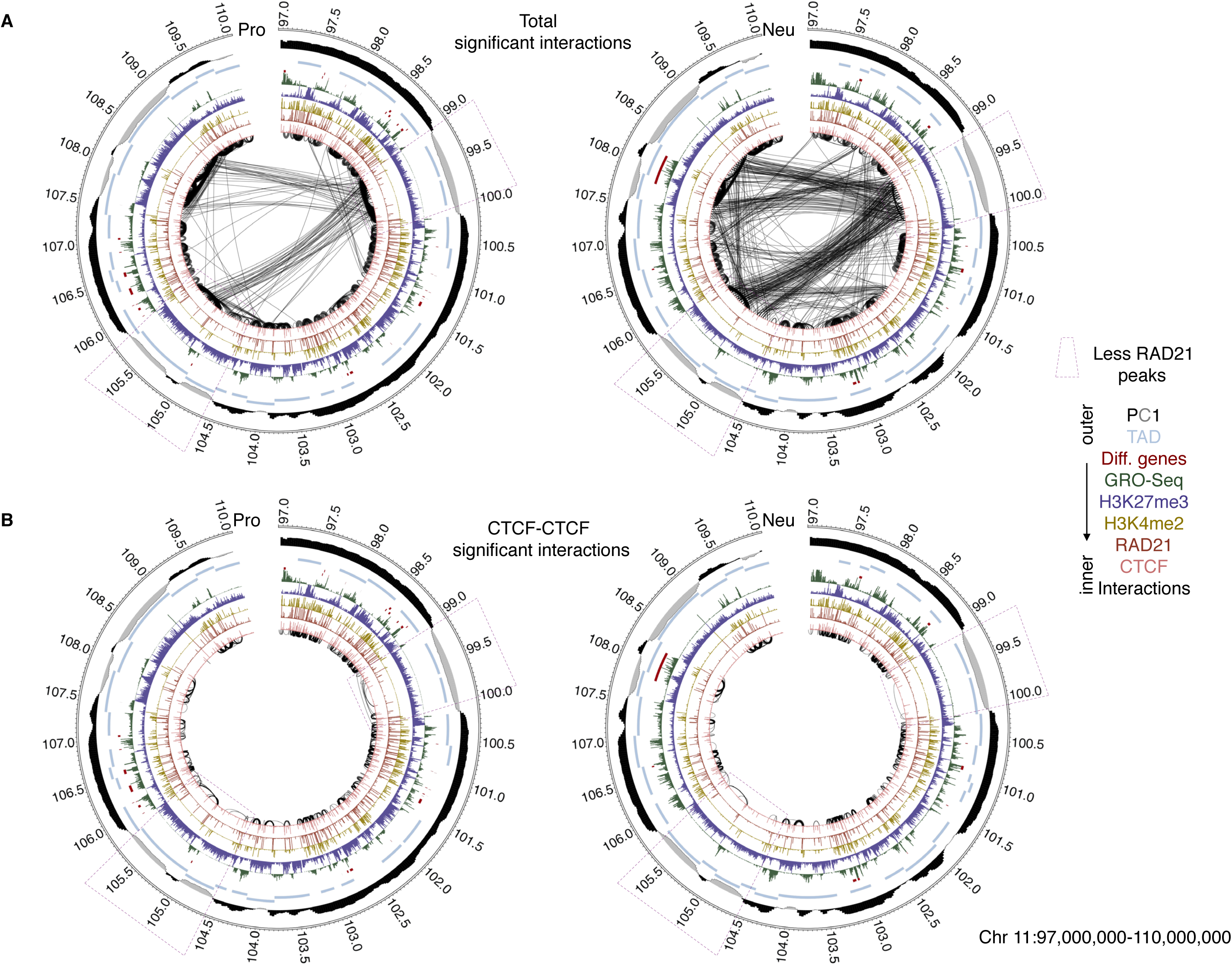
Changes in RAD21 occupancy during neutrophil differentiation are closely associated with a neutrophil specific pattern of genomic interactions. (*A*) Circos plot showing interactions from progenitors and neutrophils, RAD21 and CTCF occupancy, deposition of H3K4me2, H3K27me3, nascent transcripts abundance (GRO-Seq), cell-type specific gene, TAD and PC1 values across a 13 Mb genomic region (Chr 11:97,000,000-110,000,000). Only significant interactions are shown: *P* < 0.001, binomial test. Thickness of the connecting lines reflects the significance of the interactions (-log *P*). Bin size, 50-kb. Numbers at the margins indicate genomic positions (in Mb). (*B*) Circos plot showing CTCF versus CTCF interactions for progenitors and neutrophils spanning the same genomic region as shown in (*A*). Only significant interactions with endpoints falling into a 5 kb region surrounding CTCF peaks are shown: *P* < 0.001, binomial test. Thickness of the connecting lines reflects the significance of the interaction (-log *P*). Bin size, 5-kb. Numbers at the margins indicate genomic position (in Mb). Colors indicate CTCF, RAD21, H3K4me2, H3K27me3, GRO-Seq, cell-type specific genes, TADs and PC1 values (color key, right). Note that RAD21 depletion diminished genomic interactions between RAD21-bound sites as shown in purple dashed box. Data are from one experiment.

**Supplemental Figure S6.**
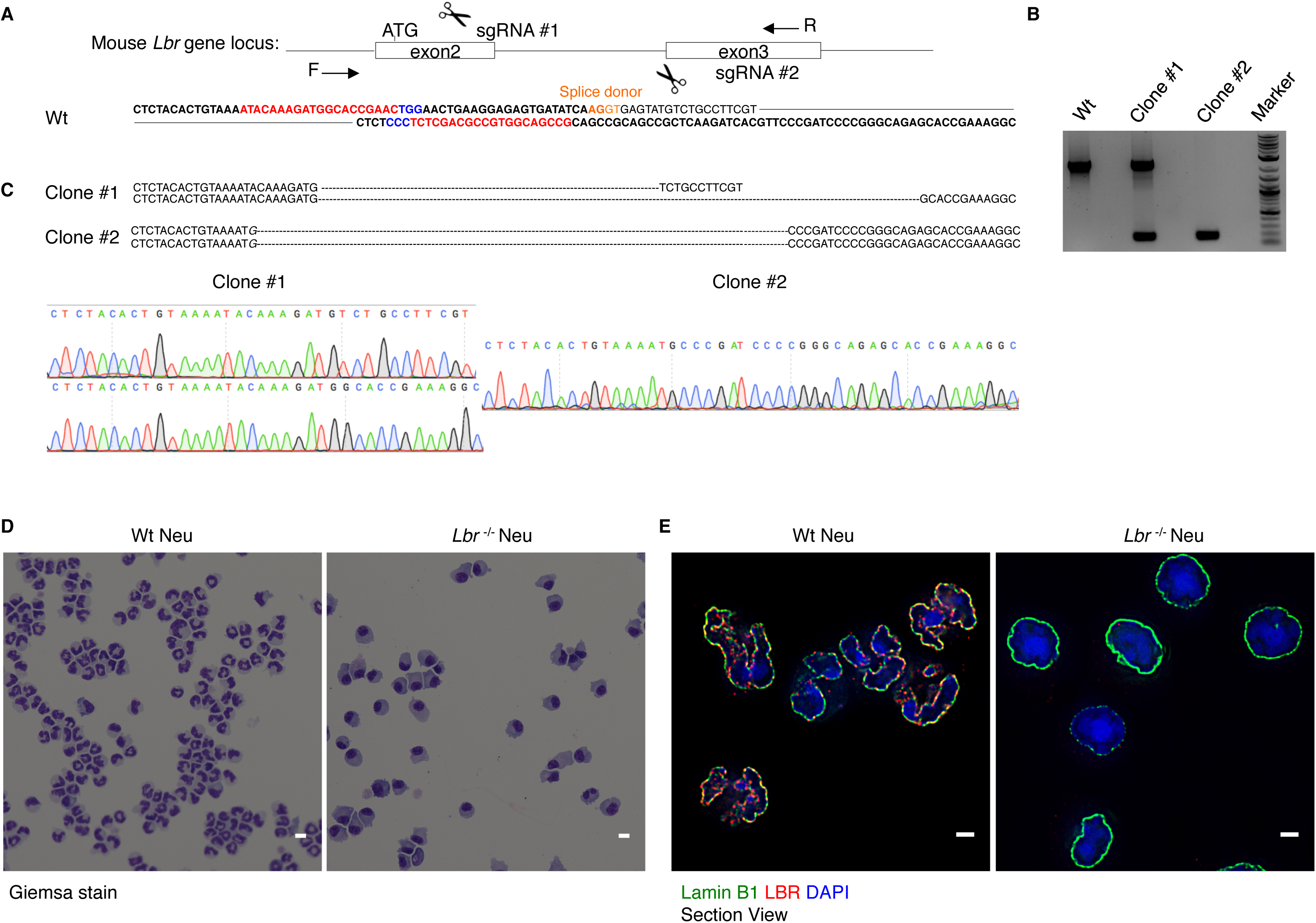
Generation of Lbr-deficient ECOMG cell lines. (*A*) Top indicates schematic diagram of Cas9/sgRNA-targeting sites close to the *Lbr* exon 2 and exon 3 junction; Bottom shows the *Lbr* genomic sequence. The exon sequences are indicated in bold. The sgRNA sequences or complimentary sequences are labeled in red. The protospacer-adjacent motif (PAM) sequences are labeled in blue. The splice donor site is labeled in orange. PCR primers (F, R) used for PCR genotyping are shown as black arrowheads. (*B*) PCR genotyping using primers F and R produced bands with correct size in targeted ECOMG clone #1 and clone #2, indicating the deletion, but not in Wt sample. (*C*) Sequences and partial chromatographs for four mutant alleles generated in two ECOMG clones. PCR products using primers F and R were sequenced. Sequence encompassing the targeted region confirmed the bi-allelic deletion. (D) Wright-Giemsa staining of Wt and *Lbr*^-/-^ neutrophils. Scale bars, 10 μm. (E) Immunofluorescence staining of Wt and *Lbr*^-/-^ neutrophils using antibodies directed against Lamin B1 and LBR. Representative image section presented as digitally magnified images. Original magnification,×100.Lamin B1, green; LBR, red; DAPI staining, blue. Scale bars, 2 μm. Note that *Lbr*^-/-^ neutrophils were associated with a substantial amount of heterochromatin that was organized into a characteristic cartwheel structure, with radial spokes extending to the periphery from a centrally located large chromocenter. LBR colocalized with Lamin B1 at the lamina in Wt neutrophils, whereas in *Lbr*^-/-^ neutrophils only background dots are detected in cytoplasm. Data are representative of three independent experiments.

**Supplemental Figure S7.**
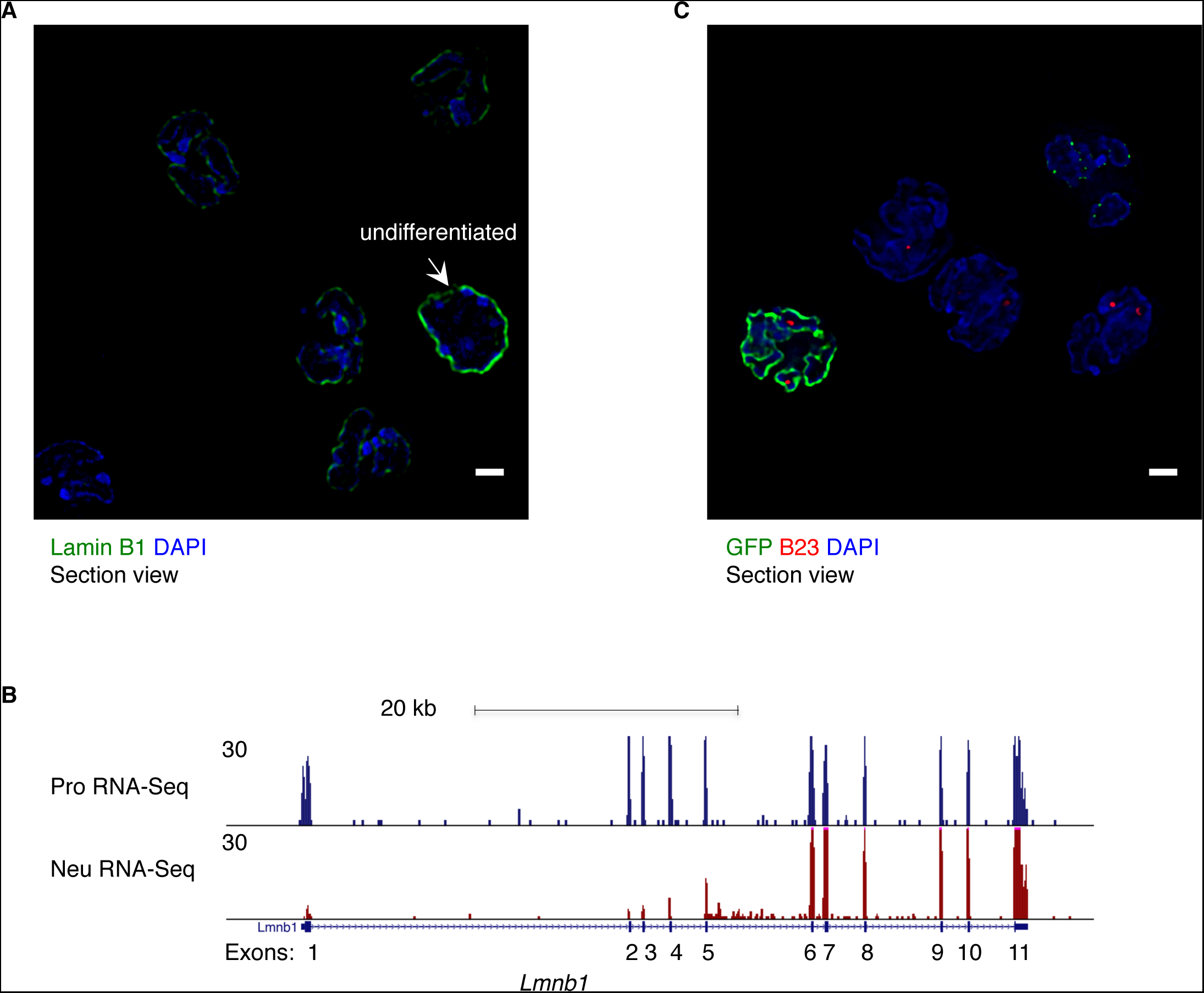
Restoring Lamin B1 expression in neutrophils. (*A*) Immunofluorescence staining in progenitors and neutrophils using a Lamin B1 antibody. Representative image section was digitally magnified. Original magnification, ×100. Lamin B1, green; DAPI staining, blue. Scale bars, 2 μm. White arrow indicates a nucleus of an undifferentiated progenitor. (*B*) Genome browser snapshot of *Lmnb1* locus showing read densities for RNA-Seq (plus strand) for progenitors and neutrophils. Note that the first five exons of *Lmnb1* were differentially spliced in neutrophils. (*C*) Immunofluorescence staining in Lamin B1-GFP transduced neutrophils using antibodies directed against GFP and B23. Representative image section presented as digitally magnified images. Original magnification, ×100. B23, red; Lamin B1-GFP, green; DAPI staining, blue. Scale bars, 2 μm. Data are representative of three independent experiments.

**Supplemental Figure S8.**
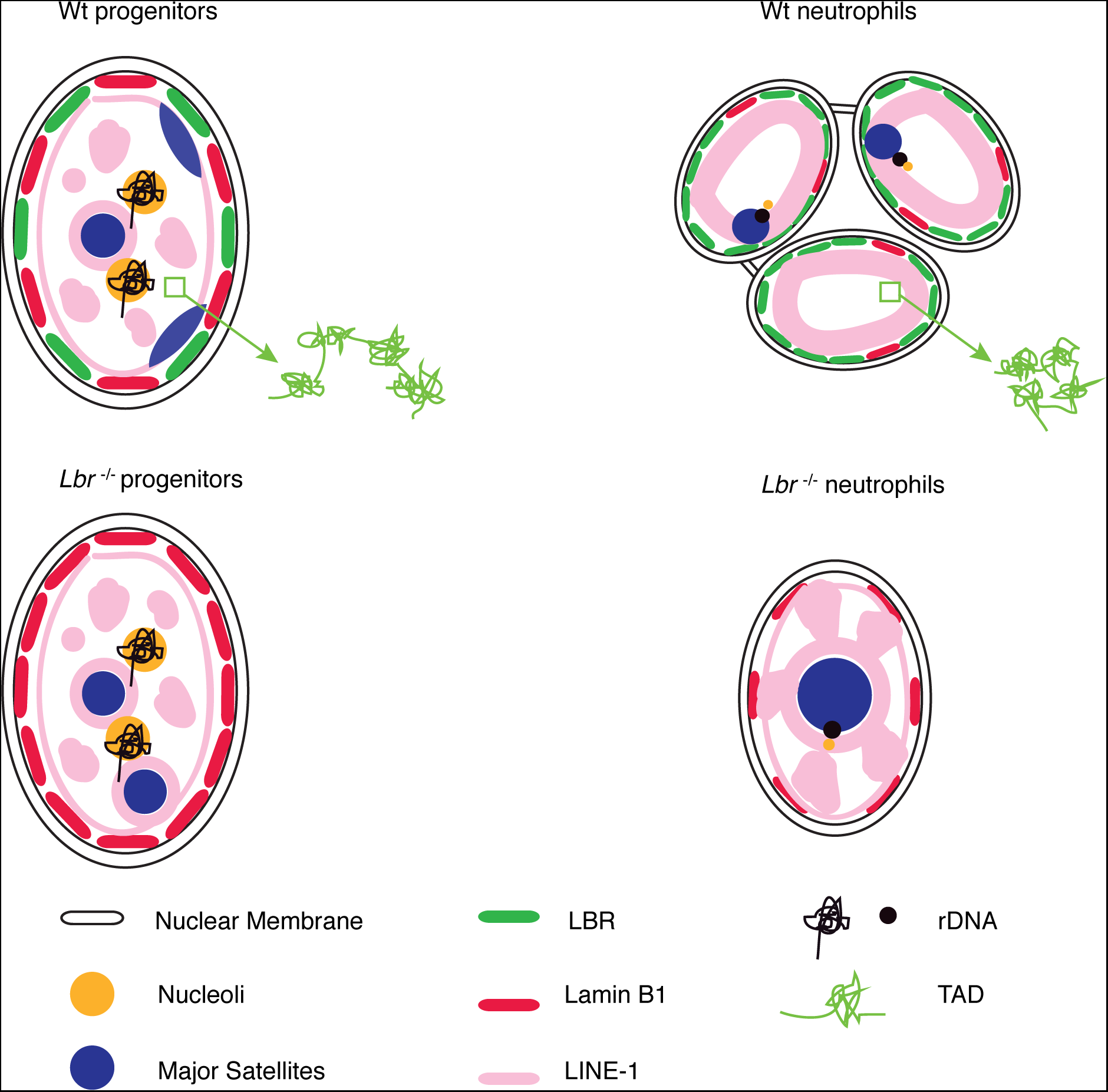
Diagram depicting the changes in genome topology associated with the transition from mononuclear to polymorphonuclear cells.

**Supplemental Table S1** List of four classes of PDs that switched compartments during neutrophil differentiation. Fold changes are shown for PDs exhibiting significant differences in nascent transcript levels (> 2-fold). Data are from one experiment.

**Supplemental Table S2** List of PDs that are most distal-favored during neutrophil differentiation is shown. Data are from one experiment.

**Supplemental Table S3** List of genes whose expression is modulated and switched compartments or changed levels of H3K27me3 during the neutrophil differentiation. Data are from one experiment.

**Supplemental Table S4** Summary of statistics of Hi-C experiments.

## Supplemental Methods

### Primary neutrophil isolation

C57BL/6J mice were housed in specific pathogen-free conditions in accordance with the Institutional Animal Care and Use Committee of University of California, San Diego. Bone marrow derived neutrophils were isolated using an EasySep™ Mouse Neutrophil Enrichment Kit (Stem Cell Technologies, 19762). Bone marrow cells were harvested from femur, tibia and crista iliac. Bone marrow cells were incubated with a cocktail of biotinylated lineage specific antibodies (EasySep™ Mouse Neutrophil Enrichment Cocktail), conjugated with EasySep™ Biotin Selection Cocktail, labeled with EasySep™ D Magnetic Particles, followed by depletion of biotin-labeled cells with EasySep™ Magnet. The enriched neutrophil purity (CD11b^+^Ly6G^+^) > 80% was validated by flow cytometry.

### Morphological evaluation

ECOMG progenitors, *in-vitro* differentiated neutrophils and BMDN were gently centrifuged (1000 rpm, 1 min) onto coverslips by Cytospin. Morphological evaluation was performed with Wright-Giemsa stain (Sigma, WS16, GS500).

### CRISPR/Cas9 mediated genome editing

Genome editing was performed using CRISPR/Cas9 essentially as described (Cong et al. 2013). Briefly, target-specific oligonucleotides were cloned into a plasmid carrying a codon-optimized version of Cas9. sgRNA sequences were cloned into the BbsI recognition sites as described (http://www.genome-engineering.org/crispr/). The sequences of guide RNAs and PCR primers are listed below. sgRNAs targeting *Lbr* exons 2 and 3 were transfected into ECOMG progenitors. Transfection was carried out with the Neon kit (Invitrogen) according to the manufacturer’s instructions. Two days after transfections, cells were plated at clonal density. Individual colonies were picked, expanded, and genotyped by PCR for deletion. The edited alleles were cloned and verified by Sanger sequencing.

sgRNA #1 top: CACCGAT ACAAAGATGGCACCGAAC

sgRNA #1 bottom: AAACGTTCG GTGCCATCTTTGTATC

sgRNA #2 top: CACCGCGGCTGCCACGGCGTCGAGA

sgRNA #2 bottom: AAACTCTCGACGCCGTGGCAGCCGC

PCR primer F: TGCCAAGTAGGAAGTTTGTTGAGG

PCR primer R: CCTCATGGGAAGCAGAGACGGATC

### RNA-Seq

ECOMG RNA was isolated and sequenced as previously described (Bossen et al. 2015). mRNA was purified from total RNA with a Dynabeads mRNA purification kit (Life Technologies). Libraries were prepared with the TruSeq primer set and were selected by size by 8% PAGE and sequenced for 50 cycles on Illumina HiSeq 2000. RNA-Seq reads were aligned to the mm9 reference genome using Tophat2, using Bowtie and Samtools. RPKMs for each RefFlat gene were calculated from aligned reads using Cufflink. For a gene to be considered expressed the cutoff of FPKM is 0.5.

### GRO-Seq

GRO-Seq was performed as previously described (Lin et al. 2012). Duplicates of GRO-Seq experiments were performed on progenitors and neutrophils. Libraries were prepared with custom GRO-seq PCR primers and were selected by size by 8% PAGE and sequenced for 50 cycles on Illumina HiSeq 2000. Reads were aligned to mm9 with Bowtie2 and reads non-uniquely mapped were discarded. Data were analyzed with HOMER (Lin et al. 2012).

### ChIP-Seq

Chromatin was immunoprecipitated as previously described (Lin et al. 2012). Antibodies used in these experiments were as follows: antibody to anti-H3K27ac (ab4729; Abcam), anti-H3K4me1 (ab8895; Abcam), anti-H3K4me2 (ab7766; Abcam), anti-H3K4me3 (ab8580; Abcam), anti-H3K27me3 (07-449; Millipore), anti-H3K9me3 (ab8898; Abcam), anti-H3K36me3 (ab9050; Abcam), anti-CTCF (07-729; Millipore) and anti-RAD21 (ab992; Abcam). Libraries were prepared with the NEBNext primer set and were selected by size by 8% PAGE and sequenced for 50 cycles on Illumina HiSeq 2000 or 2500. Reads were aligned to mm9 with Bowtie2 and reads non-uniquely mapped were discarded. Data were analyzed using HOMER. The position of each tag 3’ of its position was adjusted by 150 bp. One tag from each unique position was examined to eliminate peaks generated by clonal amplification. CTCF or RAD21 peaks were identified by searching for groups of tags located in a sliding 200-bp window. Adjacent peaks were required to be separated from each other by at least 500 bp. The threshold for the number of tags that generate a peak was selected for a FDR of 0.001. Additionally, required peaks were to have at least 4-fold more tags (normalized versus total number) than input control samples. They were also required to have 4-fold more tags relative to the local background region in a 10-kb region to avoid identification of DNA segments containing genomic duplications or nonlocalized occupancy. Peaks were associated with gene products by identification of the nearest TSS. Variable sizes of H3K27me3- marked regions were identified by searching for groups of tags located in a sliding 10-kb window. Regions were required to contain at least 1.5-fold more tags (normalized versus total number) than input control samples. Adjacent regions were required to be separated from each other by at least 50 kb. Clustering of data was done using Cluster3. Heatmaps of clusters were generated using Java Tree View.

### MeDIP-Seq

Methylated DNA IP (MeDIP) was performed as previously described (Pomraning et al. 2009) with anti-5- Methylcytosine (5-mC) (A-1014-050, epigentek) and anti-5-Hydroxymethylcytosine (5-hmC) (39770, active motif). Two replicates of experiments were performed on progenitors and neutrophils. Libraries were prepared with the NEBNext primer set and were selected by size by 8% PAGE and sequenced for 50 cycles on Illumina HiSeq 2500. Reads were aligned to mm9 with Bowtie2 and reads non-uniquely mapped were discarded. Data were analyzed using HOMER.

### In-situ Hi-C

Cells were harvested under three different biological conditions: undifferentiated Ecomg cells, *in-vitro* differentiated neutrophils, and neutrophils isolated from murine bone marrow. For *in vitro* condition, 7 independent experiments were performed starting with 5 million cells and for BMDN, 2 independent experiments were performed starting with 1 million cells. A total of 17 independent harvests were generated processed into 19 libraries (including PCR duplicates for each BMDN harvest). *In-situ* Hi-C was performed essentially as previously described (Rao et al. 2014). Libraries were prepared with the NEBNext primer set and were selected by size by 6% PAGE and sequenced for 50 cycles on Illumina HiSeq 2500. Reads were aligned to mm9 with Bowtie2 and reads non-uniquely mapped were discarded. The raw reads were filtered and analyzed using HOMER. Hi-C statistics was processed using HiCUP(Wingett et al. 2015) (Supplemental Table S4). After stringent data filtering, including removal of PCR duplicates, self-ligation, restriction ends, continuous genomic fragments and re-ligation events, a total of 243, 261 and 31 million unique pair of contacts were obtained from progenitors, neutrophils and BMDN combined replicate Hi-C libraries, respectively. The resulting neutrophil data set had more unique pair of contacts than the progenitor data set. To allow direct comparisons across progenitors and neutrophils, we resampled the neutrophil data set such that an equal number of unique pair of contacts contributed to each Hi-C data matrix. Normalized contact matrices were obtained by assuming that each genomic bin interacts with other bins with equal chances and that such interaction depends on their linear distance along the chromosome. The normalized contact probability heatmaps were generated by Java Tree View or R. PCA analysis, applied to the normalized interaction matrix, to define sub-nuclear compartments and PC1 values for each 25-kb bin with positive and negative values, corresponding to A and B compartments, respectively. We consider the difference in PC1 value (> 20) and the PC1 value change from positive to negative or vice versa to identify genomic bins that switch compartments. For data analysis in Supplemental Fig. S3B, we defined that a gene is associated with a compartment flip event if the gene promoter or gene body is overlapped with or is located within a genomic bin with such a compartment flip. Significantly interacting regions were identified as those regions that showed a markedly higher probability than expected based on a binomial test (*P* < 0.001). The circular format plots were generated using Circos. The TAD coordinates were identified by TopDom (Shin et al. 2015). Interchromosomal contact probability index (ICP) was calculated as the sum of a region's interchromosomal contact frequencies divided by the sum of its inter- and intrachromosomal contact frequencies (Kalhor et al. 2012). The distal ratio was calculated using SeqMonk (http://www.bioinformatics.bbsrc.ac.uk/proiects/seqmonk/).

### Immuno 3D-FISH and imaging

DNA-FISH was done as previously described (Lin et al. 2012). The bacterial artificial chromosome (BAC) probes were RP24-322E3, RP23-333E5, RP23-21G14, RP24-258K7, RP23-6H7, RP23-139O3, RP23-28K9, RP23-61I14, RP23-57D4, RP23-331F6, RP23-21F10, RP23-382C22, RP23-349B11 for chr17, and RP23-225M6 for rDNA (Grozdanov et al. 2003) from the BACPAC Resource Center at Children's Hospital Oakland Research Institute. Probes against mouse major satellite repeats were generated by PCR from mouse genomic DNA using the following primers: major satellite 5’- GCGAGAAAACTGAAAATCAC and 5’-TCAAGTCGTCAAGTGGATG; LINE-1 probe was generated from a cloned 3.6kb LINE-1 sequence using primers: LINE-1 5’- ACTCAAAGCGAGGCAACACTAGA and 5’- GTTCATAATGTTGTTCCACCT; Centromere probe 5’-ATTCGTTGGAAACGGGA was purchased from PNA Bio. In all cases, probes were labeled by nick-translation using Alexa-647 dUTP. The nuclear lamina was stained first with primary antibodies to Lamin B1 (sc-6217; Santa Cruz Biotechnology), followed by secondary staining with donkey antibody to goat IgG conjugated to Alexa-488 (A11055; Invitrogen). The nucleolus was stained first with primary antibodies to B23 (ab10530; Abcam), followed by secondary staining with donkey antibody to mouse IgG conjugated to Alexa-647 (A31571; Invitrogen). To check for LBR expression and localization, cells were first stained with anti-LBR (12398-1-AP; proteintech), followed by secondary staining using a donkey antibody to rabbit IgG conjugated to Alexa- 568 (A10042; Invitrogen).

RNA-FISH was performed as previously described. Cells were washed with PBS and subsequently fixed with paraformaldehyde at a final concentration of 4% for 10 minutes. After 10 minutes the cells were spun down for 2 min at 1000rpm. The fixed cells were attached to coversplis coated with 0.1% poly-l-lysine (Sigma) by cytospin. To permeabilize the cells, the cells were placed in 70% ethanol overnight. RP23-225M6 for rRNA was labeled by nick-translation using Alexa-647 dUTP, and hybridized with standard FISH hybridization buffer containing 50% formamide. For hybridization conditions 75 ng probes per 15 μ! of hybridization buffer were used. The probes were hybridized for 16 h at 42^o^C followed by two wash steps with wash buffer containing 50% formamide and 2x SSC. The cells were counterstained with DAPI before imaging.

Images were acquired on Deltavision deconvolution microscope with a 100× objective. Optical sections (z stacks) 0.2 μm apart were obtained in the DAPI, FITC, Red and Cy5 channels. Distances between FISH foci to nuclear lamina were measured with Volocity 3D image analysis software (Perkin Elmer). To measure the nuclear volume and longest axis, nuclei were counterstaining with DAPI. Images were captured using Olympus FV1000 Spectral Deconvolution Confocal microscope with a 60× objective. Using Volocity software, the volume and longest axis of DAPI stained nuclei was determined by measuring the calibrated number of voxels identified within a given nucleus.

### Statistics

*P* values of less than 0.05 were considered statistically significant. *P* values are reported in figures; the method of statistic analysis and the exact value of sample number (n) are stated in figure legends. All the tests are two-tailed and unpaired. For statistical comparison of two groups, Student’s t-test or Mann-Whitney test was performed; for multiple comparison, one-way ANOVA or Kruskal-Wallis test was performed where appropriate, using GraphPad Prism 6.0 (GraphPad Prism Software, Inc.). F test was carried to compare variabilities between the samples.

For studying the number of major satellites, centromere, rDNA foci and nucleoli, at least 100 nuclei were analyzed from each cell type. We considered that the number of observations was large enough to use Gaussian asymptotic results and then one-way ANOVA to determine significance of differences. Correct for multiple comparisons: confidence intervals and significance. Post test: Tukey. Report multiplicity adjusted *P* value for each comparison. For measuring the volume and distance from fluorescent foci to nuclear lamina, statistical analysis involved fluorescent signals obtained from > 100 cells per group. We use Kruskal-Wallis test. Corrrect for multiple comparisons: significance without confidence intervals. Post test: Dun’s. Report multiplicity adjusted *P* value for each comparison. For Hi-C interactions, the significance of the difference between observed and expected interactions was calculated using the binomial distribution based on the total number of interactions. Box and whisker plots were plotted by GrapPad Prism 6.0, where the mean value (‘‘+’’), the extreme data points (top and bottom bars), the 25^th^-75^th^ percentiles (box), and the median (Center lines) are indicated. Whisker position and outlier (dots) display are according to Tukey style.

### Publically available RNA-Seq dataset

GSE48307 (Wong et al. 2013).

